# Depletion of nuclear pore protein NUP210 suppresses metastasis through heterochromatin-mediated disruption of tumor cell mechanical response

**DOI:** 10.1101/2020.02.05.936518

**Authors:** Ruhul Amin, Anjali Shukla, Jacqueline Jufen Zhu, Sohyoung Kim, Ping Wang, Simon Zhongyuan Tian, Andy D. Tran, Debasish Paul, Steven D. Cappell, Sandra Burkett, Huaitian Liu, Maxwell P. Lee, Michael J. Kruhlak, Jennifer E. Dwyer, R. Mark Simpson, Gordon L. Hager, Yijun Ruan, Kent W. Hunter

## Abstract

Mechanical signals from the extracellular microenvironment have been implicated in tumor and metastatic progression. Here, we identified nucleoporin *NUP210* as a metastasis susceptibility gene for human estrogen receptor positive (ER+) breast cancer and a cellular mechanosensor. *Nup210* depletion suppresses lung metastasis in mouse models of breast cancer. Mechanistically, NUP210 interacts with LINC complex protein SUN2 which connects the nucleus to the cytoskeleton. In addition, the NUP210/SUN2 complex interacts with chromatin via the short isoform of BRD4 and histone H3.1/H3.2 at the nuclear periphery. In *Nup210* knockout cells, mechanosensitive genes accumulate H3K27me3 heterochromatin modification, mediated by the polycomb repressive complex 2 and differentially reposition within the nucleus. Transcriptional repression in *Nup210* knockout cells results in defective mechanotransduction and focal adhesion necessary for their metastatic capacity. Our study provides a new insight into the role of nuclear pore protein in cellular mechanosensation and metastasis.

## INTRODUCTION

The majority of cancer-related mortality is due to distant metastasis, a process in which primary tumor cells invade into surrounding endothelium, evade immunosurveillance in the circulation, travel to a distant site, extravasate, and colonize a secondary site^1–3^. This highly inefficient process requires cancer cells to employ multiple genetic and epigenetic mechanisms to establish macroscopic lesions. However, due to the limited understanding of these mechanisms, it is difficult to target metastatic cells, even with the improved therapeutic strategies^4^. Hence, improved understanding of the mechanisms which enables metastatic potential is crucial to improve patient outcomes.

Current studies have suggested that metastatic capacity in tumors may arise from the combined effect of acquired somatic mutations^5^ and epigenetic changes influencing gene expression^6, 7^. Changes in chromatin accessibility^8^ or disruptions of large heterochromatin domains^9^ can result in alteration of transcriptional programs that enable tumor cells to acquire metastatic phenotypes. In addition to these changes, previously our laboratory has demonstrated that inherited polymorphisms also have a significant effect on metastatic capacity^10^. Since the vast majority of inherited polymorphism occur in non-coding DNA, these inherited variants are thought to contribute to phenotypes by altering gene expression rather than having direct consequences on protein function. A comprehensive understanding of how the genome mediates metastatic capacity will therefore require an understanding of not only the acquired events during tumor evolution, but also how non-coding variants contribute to the complexities of gene regulation and transcriptional programs.

To gain further understanding of how polymorphisms affect regulatory elements and the subsequent metastatic phenotype, in this study we have integrated chromatin accessibility and long-range chromatin interaction analysis to identify potential metastasis susceptibility genes with polymorphic promoters and/or distant enhancers. This analysis identified *Nup210,* a gene encoding a nuclear pore complex protein, as a potential metastasis susceptibility gene. Although nuclear pore complex proteins have recently been shown to be associated with several developmental disorders and cancers^11–14^, their function in metastasis remains unexplored. Here, we established that NUP210 is responsive to mechanical signals of the extracellular microenvironment and promotes lung metastasis in mouse models of breast cancer through alteration of the mechanical response, focal adhesion, and cell migration in a nucleocytoplasmic transport-independent manner.

## RESULTS

### Identification of Nup210 as a candidate metastasis susceptibility gene

To identify polymorphic accessible chromatin regions associated with metastatic colonization, Benzonase-accessible chromatin (BACH) analysis was performed on isogenic cell lines derived from the spontaneous mammary tumor of BALB/cJ mice (67NR, 4T07 and 4T1)^15^ (Fig. 1a). When orthotopically implanted into mice, 67NR forms tumors but remains localized, 4T07 cells disseminate to distant sites but rarely form macroscopic lesions, while 4T1 cells complete the metastatic process to form multiple pulmonary macroscopic lesions.

**Figure 1:**
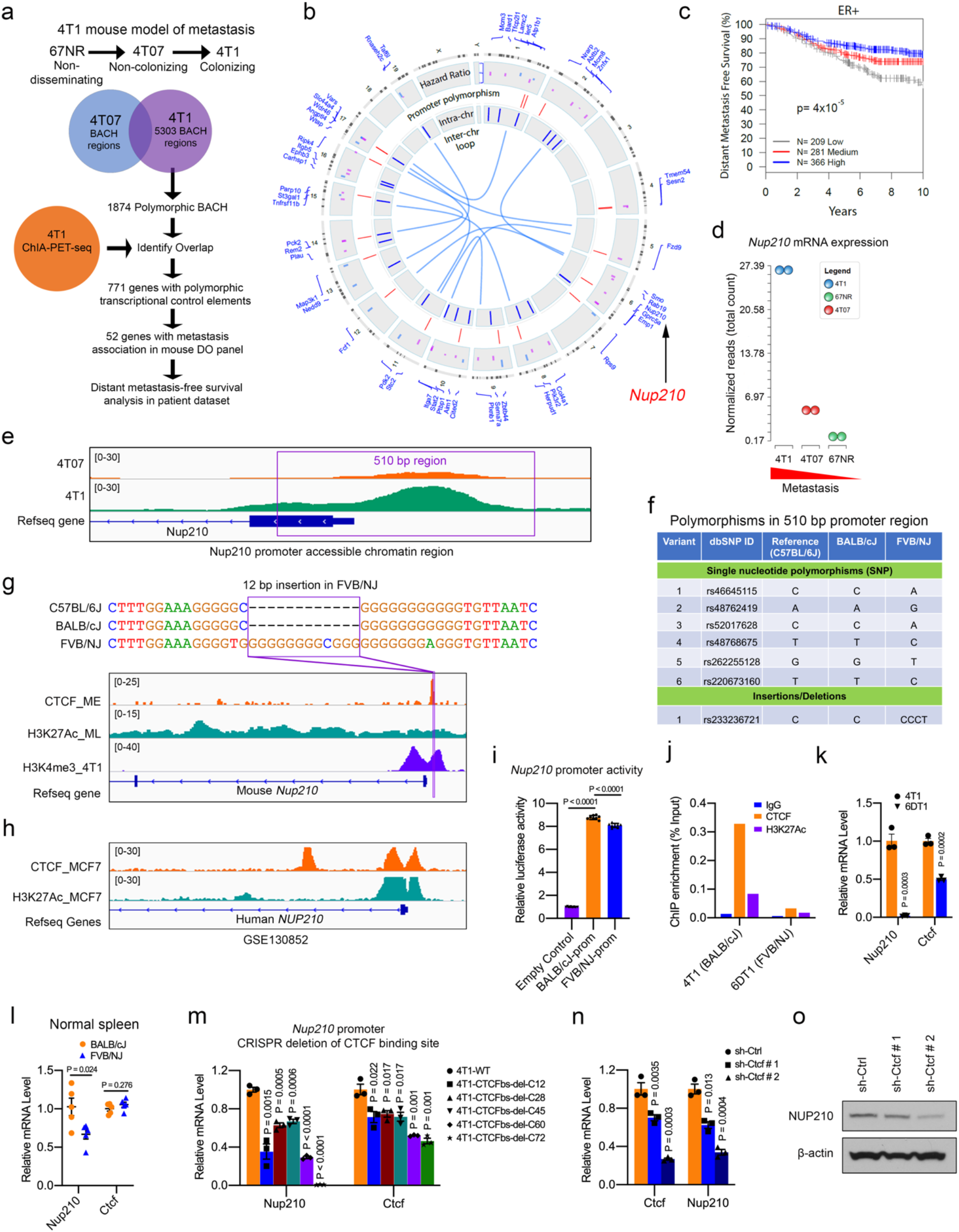
Genome-wide analysis of non-coding region polymorphisms identifies *Nup210* as a metastasis susceptibility gene. (a) Scheme for metastasis susceptibility gene identification. (b) Circos plot representation of the result in (a). Red lines = polymorphic promoter, blue lines = intra-chromosomal and light blue lines = inter-chromosomal looping interactions. (c) Distant metastasis-free survival (DMFS) of human ER+ breast cancer patients stratified by their expression of the 52-gene signature. (d) *Nup210* mRNA expression in the mouse 4T1 cell line series. (e) Integrative Genomics Viewer (IGV) track of the *Nup210* promoter BACH region in 4T07 and 4T1 cells. (f) Polymorphisms within the 510 bp *Nup210* promoter region. (g) 12 bp FVB/NJ promoter indel is located within a CTCF binding site in mouse *Nup210* promoter. (h) CTCF and H3K27Ac enrichment in human *NUP210* promoter of MCF7 cells (i) Luciferase assay of BALB/cJ and FVB/NJ *Nup210* promoter regions. ANOVA, tukey’s multiple comparison test, mean ± s.e.m. (j) ChIP analysis of CTCF and H3K27Ac at Nup210 promoter of 4T1 and 6DT1 cells. (k) *Nup210* and *Ctcf* mRNA levels in cell lines (4T1, 6DT1) and (l) mouse spleen (BALB/cJ or FVB/NJ). (m) *Nup210* and *Ctcf* mRNA level in *Nup210* promoter CTCF binding site-deleted clones. Two tailed t-test, mean ± s.e.m. (n) Effect of *Ctcf* knockdown on 4T1 *Nup210* mRNA and (o) protein levels. Two tailed t-test, mean ± s.e.m.

To identify transcriptional control regions enriched for the dormant-to-proliferative switch during metastatic colonization, 5303 BACH sites unique to the 4T1 cell line were identified. These sites were then intersected with the polymorphisms present in the eight founder strains^16^ of the mouse Diversity Outbred (DO) genetic mapping panel^17^. Since the vast majority of these sites did not fall within the proximal promoters of protein coding genes, Chromatin Interaction Analysis Through Paired-end Tag Sequencing (ChIA PET-seq) was performed in 4T1 cells to putatively link polymorphic transcriptional control elements (promoters and/or enhancers). Finally, the resulting gene list was filtered through expression data of mice resulting from a cross between the MMTV-PyMT model^18^ and the DO to identify genes associated with metastasis in this population. This strategy identified 52 potential metastasis susceptibility genes (Fig. 1b). To assess human relevance, the hazard ratios calculated for these genes in the mouse data were used to generate a weighted gene signature that was subsequently screened on human breast cancer gene expression datasets. Significant stratification of distant metastasis-free survival (DMFS) in estrogen receptor-positive (ER+) breast cancer was observed (Fig. 1c), consistent with a potential role of these genes with human breast cancer progression. NUP210, a nuclear pore protein, was selected for further validation due to the association of a number of previously identified metastasis susceptibility genes with the nuclear envelope^19^. To test the function of *Nup210* in metastasis, we initially asked whether variation in *Nup210* expression would correlate with metastatic potential. Our RNA-seq data from the 4T1 series of cell lines revealed that *Nup210* expression is higher in metastatic than non-metastatic variant cell lines (Fig. 1d) suggesting a positive correlation of Nup210 expression and metastatic potential.

### Polymorphisms in the Nup210 promoter affect CTCF binding and Nup210 transcription

To examine the causal variant responsible for the transcriptional difference in the mouse genetic screen, the proximal promoter of *Nup210* was identified based on the BACH profile (Fig. 1e). Analysis of the Mouse Genomes Project database^16^ revealed 6 single nucleotide polymorphisms (SNPs) and one insertion/deletion (indel) in a 510 bp promoter region of *Nup210* (Fig. 1f). This included a 12 bp G-rich insertion in the *Nup210* promoter of the FVB/NJ, the MMTV-PyMT parental strain (Fig. 1g), located within a CTCF binding region enriched in enhancer (H3K27Ac)-promoter (H3K4me3) marks^20^. CTCF binding on Nup210 promoter also appeared to be evolutionarily conserved between mouse and human based on analysis in MCF7 human breast cancer cell line (Fig. 1h). We initially tested whether the polymorphisms within *Nup210* promoters can affect gene expression via cloning these 510 bp regions into a promoter luciferase reporter. A moderate (∼10%), but significant, reduction of luciferase activity was observed for the FVB/NJ-derived *Nup210* promoter region compared to the BALB/cJ promoter indicating a functional impact of the polymorphism on *Nup210* transcription (Fig. 1i). Then we examined the effect of 12bp indel on CTCF binding and *Nup210* transcription as CTCF is known to regulate gene expression^21, 22^. CTCF was preferentially enriched in the *Nup210* promoter region of 4T1 cells (derived from BALB/cJ mice) compared with 6DT1 cells (derived from FVB/NJ mice) (Fig. 1j). Consistently, expression of *Nup210* and *Ctcf* were significantly higher in 4T1 cells than in 6DT1 cells (Fig. 1k). In addition, *Nup210* expression was significantly lower in the normal spleen of FVB/NJ compared to BALB/cJ mice (Fig. 1l) despite similar levels of CTCF. CRISPR/Cas9-mediated deletion of CTCF binding site on 4T1 *Nup210* polymorphic promoter showed significantly decreased *Nup210* mRNA which further supported that *Nup210* promoter indel is a direct target of CTCF (Fig. 1m, Extended Data Fig. 1a-c). *Ctcf* mRNA was also downregulated in these deletion clones indicating a potential positive feedback loop between NUP210 and CTCF. Furthermore, *Ctcf* knockdown in 4T1 cells revealed a marked reduction of *Nup210* both at the transcript (Fig. 1n) and protein level (Fig. 1o) suggesting a direct effect of CTCF loss on *Nup210* transcription. Taken together, these results suggest that *Nup210* is a CTCF-regulated gene and the indel in the promoter alters *Nup210* transcription by modulating CTCF binding to the promoter region.

### NUP210 expression is associated with metastasis in human breast cancer patients

Consistent with the mouse data, *NUP210* amplification in human breast cancer patients was found to be associated with decreased overall survival in the METABRIC dataset^23^ (Fig. 2a). Two independent human breast cancer gene expression datasets suggested that *NUP210* mRNA expression was significantly associated with reduced overall survival (Fig. 2b, METABRIC) and distant metastasis-free survival (DMFS; Fig. 2c, Km-plotter)^24^ in ER+ breast cancer patients. However, the patient survival outcome appears to be moderate in these datasets suggesting that NUP210 expression is likely one of the multiple factors contributing to the patient outcome. Although association of *NUP210* mRNA expression among triple-negative (ER-/PR-/HER2-) breast cancer patients was not consistently significant in these two datasets (Extended Data Fig. 2a, b), recent proteomics analysis revealed that higher NUP210 protein level is significantly associated with the poor DMFS of triple-negative cancer patient (Km-plotter) (Extended Data Fig. 2c)^25^. METABRIC data suggests that *NUP210* expression is higher in luminal B and basal subtypes than in luminal A, Her2+, and claudin-low subtypes of breast cancer (Fig. 2d). Consistently, immunohistochemical analysis in human normal mammary gland revealed that NUP210 is predominantly expressed in luminal cell compartment (Fig. 2e). Analysis of a small human breast cancer tissue microarray revealed that NUP210 protein level was heterogenous among the primary tumors of both ER+ and ER- patients (Fig. 2f). Interestingly, in addition to nuclear envelope staining of NUP210, cytoplasmic staining was also observed in many cases suggesting that NUP210 might be mislocalized in some patients. Confining the quantification of NUP210 signal intensity at the nuclear envelope revealed no significant difference between ER+ and ER- primary tumors (Fig. 2g). However, NUP210 level was significantly higher in the lymph node metastases of ER+ than ER- patient (Fig. 2h, i), consistent with the survial outcome of breast cancer patient in different datasets. Analysis in publicly available gene expression data of breast cancer patients revealed that visceral metastases (lung, liver) have significantly higher expression of *NUP210* than nonvisceral metastases (lymph node) (Fig. 2j). Furthermore, *NUP210* expression is significantly higher in metastases than primary tumor of prostate cancer (Extended Data Fig. 2d) and melanoma (Extended Data Fig. 2e). These data indicate that *NUP210* is associated with human cancer progression and further supports its potential role as a metastasis susceptibility gene.

**Figure 2:**
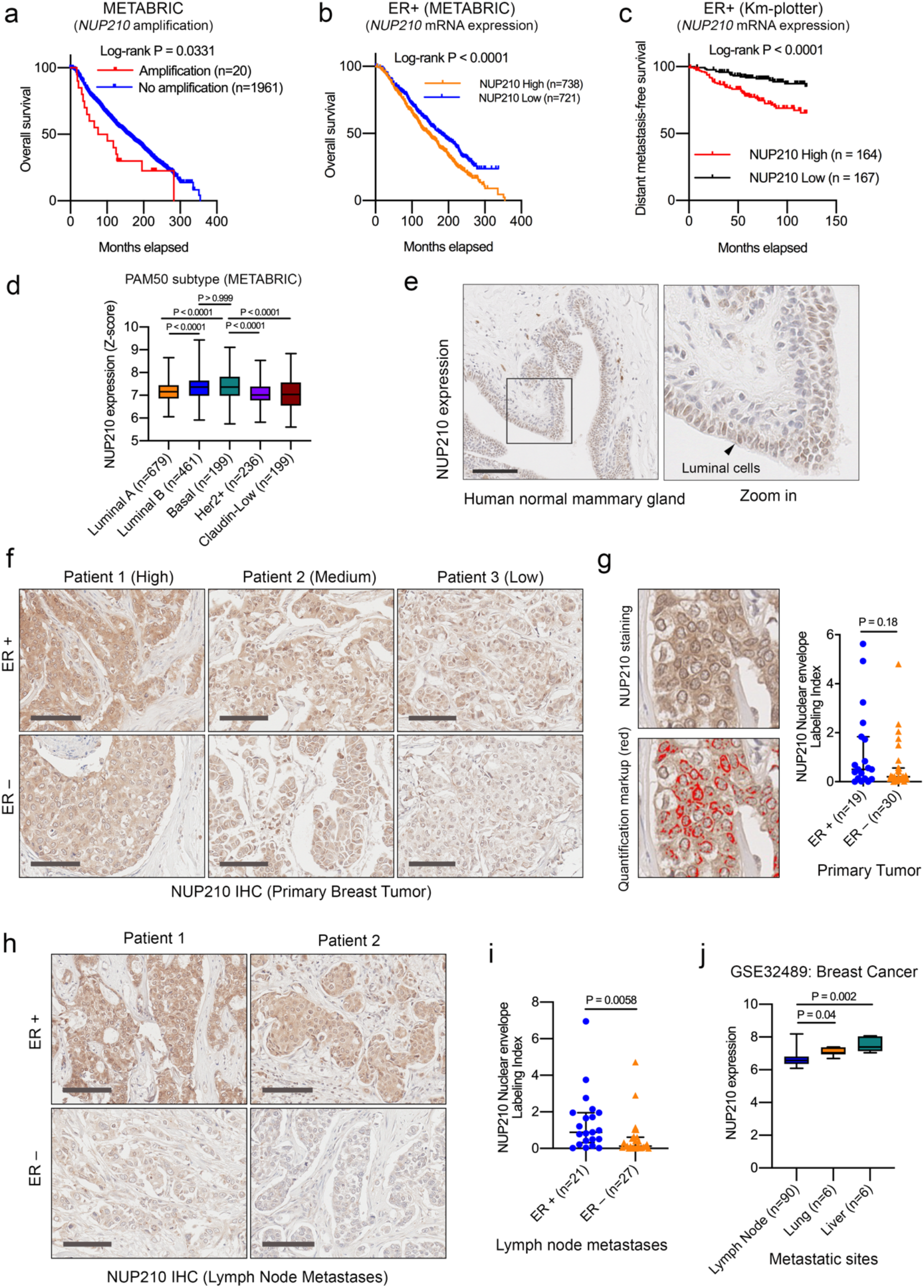
*NUP210* expression is associated with poor outcome in human ER+ breast cancer patients. (a) Association of *NUP210* amplification on the overall survival (OS). (b) Association of *NUP210* mRNA level on the OS and (c) DMFS in ER+ patients. (d) *NUP210* mRNA level within METABRIC dataset. Kruskal-Wallis ANOVA test. (e) NUP210 protein expression in human normal mammary gland. Scale bar 100 μm. (f) Immunohistochemistry and nuclear envelope quantification (g) of NUP210 in the primary tumors of ER+ and ER- cancer patients. Scale bar 100 μm. Mann-Whitney U test. (h) Immunohistochemistry analysis and nuclear envelope quantification (i) NUP210 protein in the lymph node metastases of ER+ and ER- patients. Scale bar 100 μm. Mann-Whitney U test. (j) *NUP210* mRNA level in multiple human breast cancer metastatic sites. Kruskal-Wallis ANOVA test.

### Depletion of Nup210 in cancer cells decreases lung metastasis in mice

To validate the role of *Nup210* in metastasis, luminal-like orthotopic mammary tumor transplantation models were used^26^. 3 different cell lines were utilized for this analysis: 4T1^15^, 6DT1, derived from the mammary tumor of FVB/MMTV-MYC transgenic mouse, and MVT1, derived from the mammary tumor of FVB/MMTV-MYC/VEGF double transgenic mouse^27^. Primary tumors derived from orthotopically implanted *Nup210* shRNA knockdown (Fig. 3a, Extended Data Fig. 3a) cells showed variability in tumor weight, with decreased tumor weight for 4T1 cells (Fig. 3b) but increased tumor weights for both 6DT1 and MVT1 cells (Extended Data Fig. 3b). However, *Nup210* knockdown resulted in a decrease in pulmonary metastases for all three cell lines (Fig. 3c, d; Extended Data Fig. 3c-f), and this decrease was preserved after normalizing metastasis counts by tumor weight to account for the variability in primary tumor growth. These results were therefore consistent with the association of *NUP210* with poor prognosis in patients with ER+ breast cancers.

**Figure 3:**
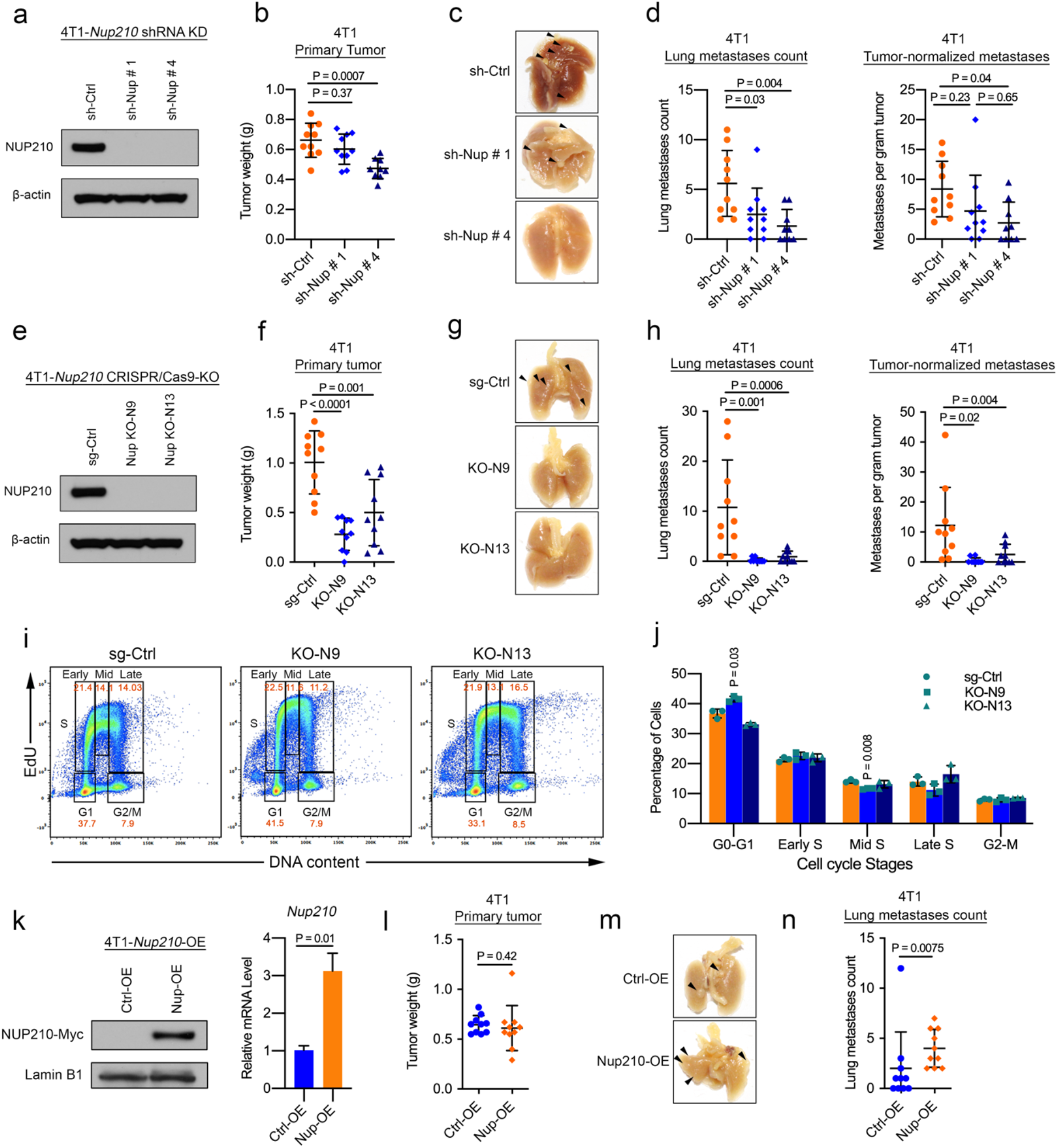
Depletion of *Nup210* in 4T1 metastatic cancer cells reduces lung metastasis in mice. (a) Western blot of *Nup210* knockdown (KD) in 4T1 cells. (b) Primary tumor weight, (c) representative lung images, (d) lung metastases count and lung metastases count normalized to primary tumor weight in orthotopically transplanted *Nup210* KD 4T1 cells. ANOVA, Tukey’s multiple comparison test, Mean ± S.D. (e) Western blot showing CRISPR/Cas9-mediated knockout (KO) of *Nup210* in 4T1 cells. (f) Primary tumor weight, (g) representative lung images, (h) lung metastases count and tumor-normalized metastases count after orthotopic transplantation of *Nup210* KO 4T1 cells. ANOVA, Tukey’s multiple comparison test, mean ± s.d. (i) DNA content analysis with EdU incorporation of the different stages of the cell cycle in sg-Ctrl and *Nup210* KO 4T1 cells. (j) Quantification of cell cycle stage distribution in sg-Ctrl and *Nup210* KO 4T1 cells.Two tailed t-test, mean ± s.e.m, n=3. (k) Western blot (left) and qRT-PCR (right) of NUP210 overexpression in 4T1 cells. Two tailed t-test, mean ± s.e.m, n=3. (l) Primary tumor weight, (m) representative lung images, (n) lung metastases count after orthotopic transplantation of NUP210-overexpressing 4T1 cells. Mann Whitney-U test, mean ± s.d.

To further validate the role of *Nup210* in metastatic progression, CRISPR/Cas9-mediated knockout (KO) of *Nup210* was performed in the 4T1 cell line (Fig. 3e). Similar to shRNA result, *Nup210* KO in two different clones, N9 and N13, diminished primary tumor weight and lung metastases (Fig. 3f-h). Although we observed differential effects of *Nup210* loss on primary tumor weight, cell cycle analysis on *Nup210* KO 4T1 cells (Fig. 3i, j) revealed no consistent changes in the distribution of cell population among cell cycle stages, suggesting that the effect of NUP210 loss on tumor and metastasis is independent of a cell-intrinsic ability of NUP210 to regulate proliferation. Finally, 4T1 cells with overexpression of *Nup210* showed no difference in primary tumor weight (Fig. 3k, l) but lung metastasis was significantly increased (Fig. m, n). Taken together, these results indicate that *Nup210* promotes lung metastasis in mouse models of luminal breast cancer.

### NUP210 tethers histone H3.1/3.2 to the nuclear periphery

To investigate the mechanism by which NUP210 alters metastatic function, initially we examined the effect of NUP210 depletion on nucleocytoplasmic transport, a general function of nuclear pore. A recent study has suggested that loss of NUP210 does not affect the general nucleocytoplasmic transport in differentiating myotubes^28^. Consistent with this, transfection of NES-tdTomato-NLS nucleocytoplasmic transport reporter (Extended Data Fig. 4a) into *Nup210* knockdown 4T1 cells with or without the nuclear export inhibitor leptomycin B showed no significant differences in nucleocytoplasmic transport of the reporter protein (Extended Data Fig. 4b).

To better understand how NUP210 promotes metastasis, NUP210 co-immunoprecipitation (Co-IP)-mass spectrometry analysis was performed (Fig. 4a). Both endogenous and Myc-tagged NUP210 potentially interact with multiple chromatin-associated molecules including histone H3.1 (Fig. 4b). We decided to examine NUP210-histone H3.1 interaction due to the potential association of H3.1 with poor outcome in breast cancer patients (P = 0.0642) in METABRIC dataset (Fig. 4c) and reported mutations of histone H3.1 in human cancer^29, 30^. Reciprocal Co-IP in 4T1 cells using an antibody that recognizes both H3.1 and H3.2 validated the predicted H3.1-NUP210 interaction (Fig. 4d, e). H3.1/3.2 also pulled down lamin B1, a component of the nuclear lamina, suggesting that this interaction occurs at the nuclear periphery. Reciprocal co-IP using both endogenous and Flag-tagged H3.1 in human 293FT cells also showed the similar result (Fig. 4f). Immunofluorescence analysis confirmed the association of H3.1/3.2 and NUP210 at the nuclear periphery (Fig. 4g). Extending the analysis in human breast cancer cells revealed that NUP210-H3.1/3.2 interaction is more prominent in ER+ MCF7 cells than ER- MDA-MB-231 cells (Extended Data Fig. 5a, b). These results establish an evolutionary conserved association of NUP210 with H3.1/3.2 in mouse and human cells.

**Figure 4:**
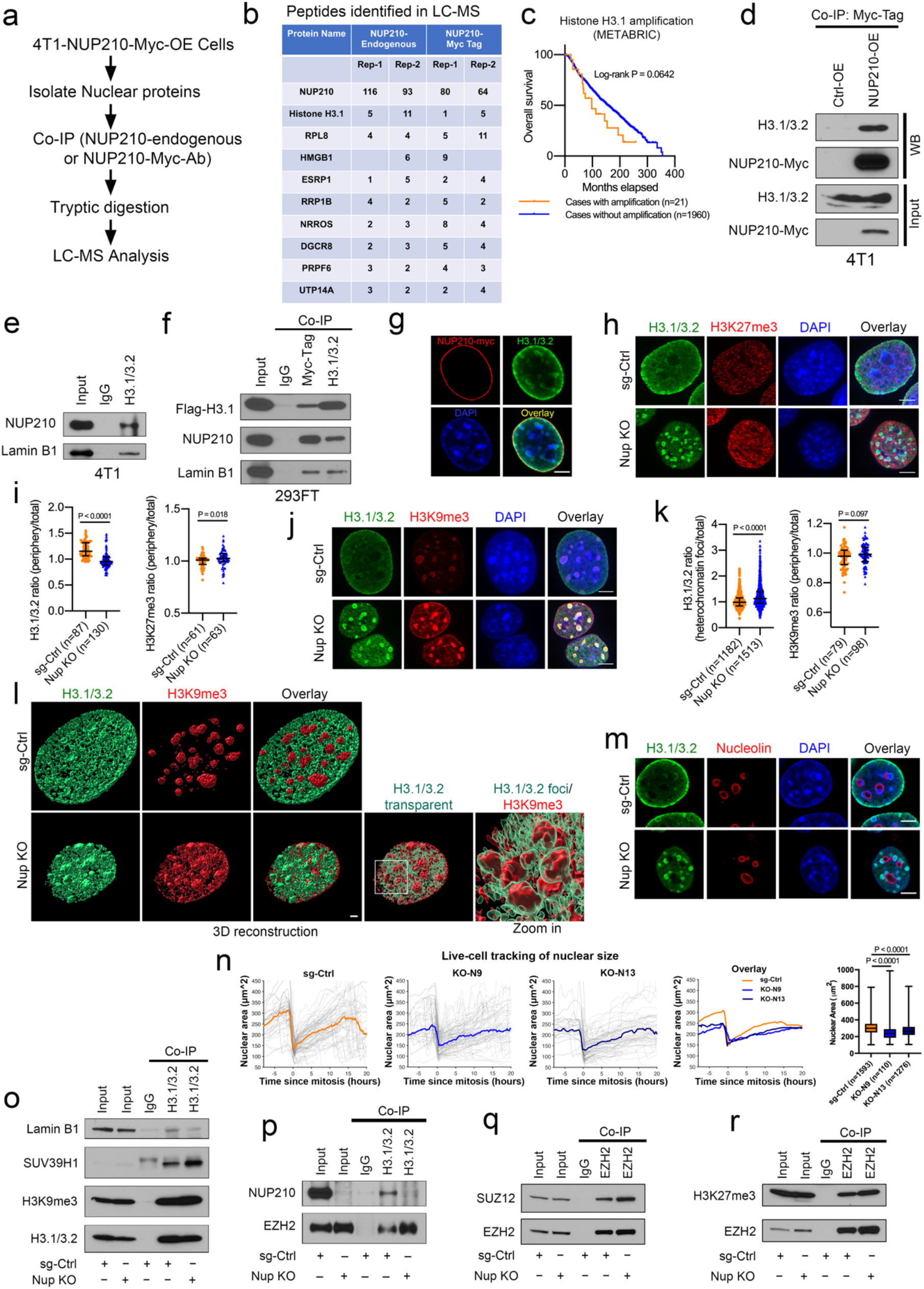
NUP210 interacts with histone H3.1/3.2 at the nuclear periphery and H3.1/3.2 is re-distributed to heterochromatin foci upon NUP210 loss. (a) NUP210 co-immunoprecipitation (Co-IP)-LC-MS analysis. (b) Peptide count in LC-MS analysis by endogenous and Myc-tag-NUP210 antibody. (c) Association of Histone H3.1 amplification on the overall survival of breast cancer patients. (d) Co-IP validation of NUP210-Myc interaction with H3.1/3.2 in 4T1 cells. (e) Co-IP validation of endogenous H3.1/3.2, NUP210, Lamin B1 interaction in 4T1 cells. (f) Reciprocal Co-IP of Flag-H3.1 and NUP210-Myc in human 293FT cells. (g) Representative immunofluorescence images of of H3.1/3.2 and NUP210-Myc at the nuclear periphery in 4T1 cells. Scale bar = 5 μm. (h) Representative images of the distribution of H3.1/3.2 and H3K27me3 in sg-Ctrl and *Nup210* KO 4T1 cells. Scale bar = 5 μm. (i) H3.1/3.2 and H3K27me3 intensity ratio (nuclear periphery vs total nuclear intensity) in *Nup210* KO 4T1 cells. Mann-Whitney U test, median with interquartile range. (j) Representative images of H3.1/3.2 localization with pericentric heterochromatin (H3K9me3) in *Nup210* KO 4T1 cells. Scale bar = 5 μm. (k) (Left) H3.1/3.2 intensity ratio (heterochromatin foci vs total nucleus) and (Right) H3K9me3 intensity ratio (periphery vs total nuclear intensity) in *Nup210* KO 4T1 cells. Mann-Whitney U test, median with interquartile range. (l) 3D reconstruction of H3.1/3.2 and H3K9me3 distribution in *Nup210* KO 4T1 cells. Display intensity was adjusted to visualize distinct foci. Scale bar = 2 μm. (m) Representative images of H3.1/3.2 distribution at the nuclolear (nucleolin) periphery in *Nup210* KO 4T1 cells. Scale bar = 5 μm. (n) Live cell imaging of the nuclear size of *Nup210* KO 4T1 cells before and after mitosis. Thin line, single nuclear size traces; thick line, median of nuclear size (n > 100 cells); box plot represents average nuclear size. Kruskal-Wallis ANOVA with Dunn’s multiple comparison test. The box extends from 25^th^ to 75^th^ percentile, whiskers extend from smallest values to the largest values, horizontal line represents median. (o) Co-IP of H3.1/3.2 with lamin B1, SUV39H1, and H3K9me3 in *Nup210* KO 4T1 cells. (p) Co-IP of H3.1/3.2 with EZH2; (q) EZH2 and SUZ12; (r) EZH2 and H3K27me3 in *Nup210* KO 4T1 cells.

### Increased recruitment of heterochromatin-modifying enzymes, SUV39H1 and EZH2, to H3.1/3.2 in Nup210 KO cells

H3.1 is known to be incorporated into nucleosomes in S phase of the cell cycle in a replication-dependent manner^31^, which suggested that NUP210 loss might affect S phase progression. As mentioned earlier, there was no consistent changes in S phase of the cell cycle in *Nup210* KO cells (Fig. 3i, j), suggesting that NUP210-H3.1/3.2 association might have a different role in metastatic cells. The effect of NUP210 loss on H3.1/3.2 interactions in metastatic cells was therefore examined. Loss of NUP210 resulted in a redistribution of H3.1/3.2 from the nuclear periphery to foci-like structures inside the nucleus (Fig. 4h, i), suggesting that NUP210 might be tethering H3.1/3.2 to the nuclear envelope. A significant enrichment of H3K27me3 heterochromatin mark at the nuclear periphery was also observed. Co-staining with H3K9me3, a pericentric heterochromatin-associated histone marker, and DAPI revealed that the redistributed H3.1/3.2 colocalized with heterochromatin in *Nup210* KO cells (Fig. 4j, k). 3D reconstruction revealed that H3.1/3.2 foci were localized on the periphery of H3K9me3-marked region (Fig. 4l, Supplementary Video 1, 2). Many of the foci appeared on the heterochromatin-enriched nucleolar periphery (Fig. 4m). Like 4T1, H3.1/3.2 was also redistributed from the nuclear periphery to intra-nuclear foci in *NUP210*-depleted ER+ MCF7 cells (Extended Data Fig. 5c, d). Although there was significant loss of H3.1/3.2 from the nuclear periphery of *NUP210*-depleted, ER-, MDA-MB-231 cells (Extended Data Fig. 5e, f), H3.1/3.2 foci was not prominent in these cells suggesting a subtype-specific effect of NUP210 loss. Live-cell imaging demonstrated that nuclear size was significantly decreased in *Nup210* KO cells throughout the cell cycle (Fig. 4n) suggesting that the loss of NUP210 might increase heterochromatinization^32^. Co-IP in *Nup210* KO cells revealed a decreased association of H3.1/3.2 with lamin B1 and increased association with heterochromatin modifying enzymes, SUV39H1 (catalyzing H3K9me3) (Fig. 4o) and EZH2 (Fig. 4p), a key component of the polycomb repressive complex 2 (PRC2) (catalyzing H3K27me3)^33^. Further analysis of other PRC2 components revealed an increased association of EZH2 with SUZ12 (Fig. 4q) and H3K27me3 (Fig. 4r) in *Nup210* KO cells. Taken together, these results suggest that NUP210 restricts the recruitment of heterochromatin-modifying enzymes to H3.1/3.2 at the nuclear periphery.

### NUP210 loss increases heterochromatin spreading and decreases expression of cell adhesion/migration-related genes

To test whether NUP210 loss promotes heterochromatin formation, we performed ChIP-seq analysis of NUP210, H3K27me3 (repressive) and H3K4me3 (active) histone marks. ChIP-seq analysis revealed that the majority (55.33%) of the NUP210 peaks are within intergenic regions (Extended Data Fig. 6a), with 19.01% of NUP210 peaks within 10 kb of transcription start sites (TSS). The rest of the peaks were found within gene bodies but mainly enriched in 3*′*-transcription end sites (TES) of the genes (Extended Data Fig. 6b). Moreover, 40% (9,962 peaks) of NUP210 peaks overlap with H3K27me3 (FDR < 0.05) (Extended Data Fig. 6c) indicating H3K27me3 enrichment surrounding the NUP210-bound gene bodies. In addition, in *Nup210* KO cells, increased H3K27me3 over NUP210-bound gene bodies was observed (Fig. 5a), suggesting that these genes were undergoing heterochromatinization. There was overall expansion of H3K27me3 peak regions, suggesting H3K27me3-marked heterochromatin spread (Fig. 5b). Integration of ChIP-seq dataset for active enhancer (H3K27Ac)^34^ revealed that NUP210 was enriched at regions surrounding H3K27Ac (Extended Data Fig. 6d). Loss of NUP210 resulted in an increase in the repressive H3K27me3 mark across these enhancer regions, suggesting NUP210 might be preventing heterochromatin spreading across distal regulator elements as well as gene bodies.

**Figure 5:**
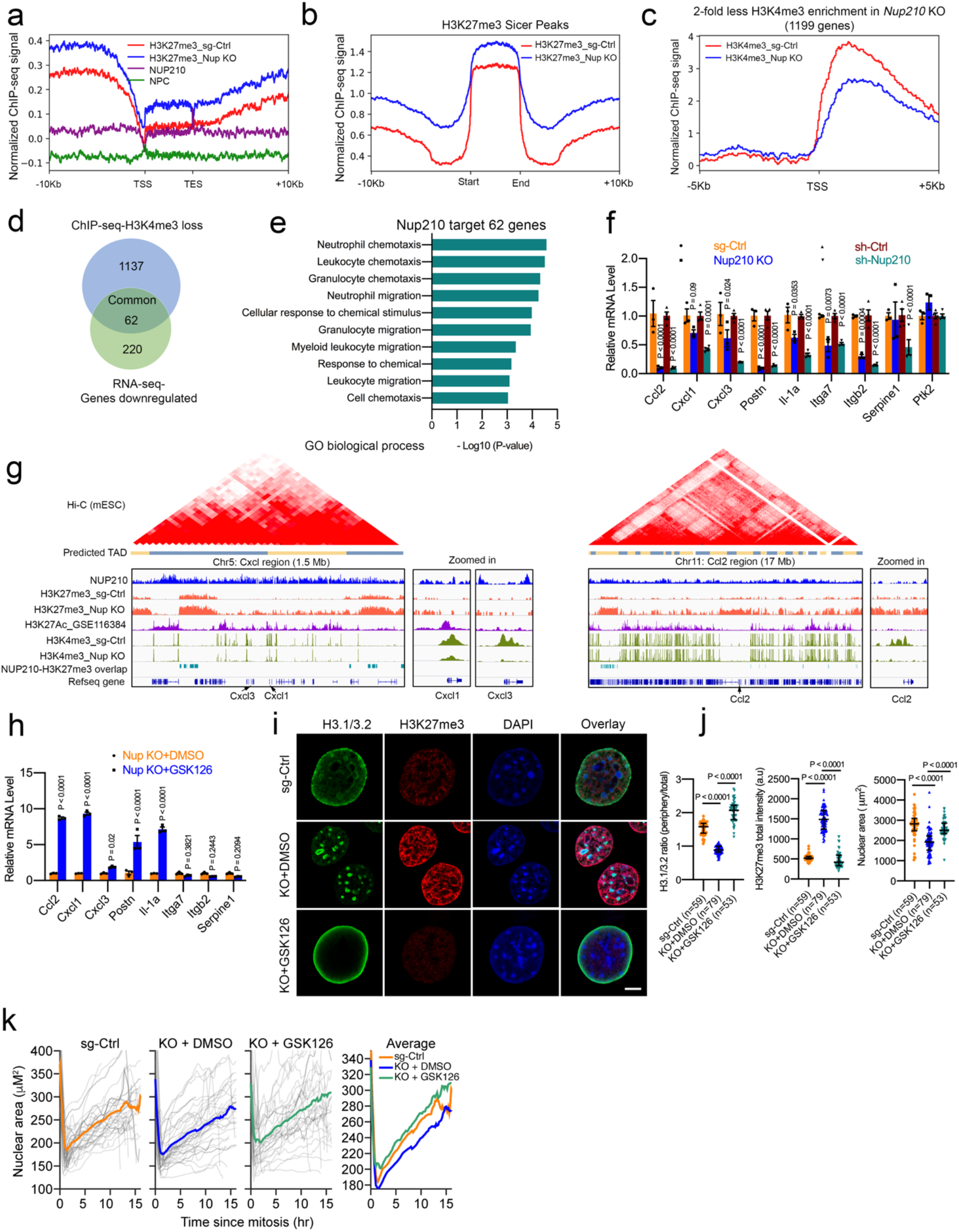
Loss of NUP210 is associated with H3K27me3-marked heterochromatin spreading in 4T1 cells. (a) H3K27me3 peak enrichment on gene bodies in *Nup210* KO cells within NUP210-enriched regions. (b) H3K27me3 ChIP-seq profile in sg-Ctrl and *Nup210* KO cells. (c) H3K4me3 ChIP-seq profile on gene promoters in sg-Ctrl and *Nup210* KO cells. (d) Overlap of genes with H3K4me3 loss on promoter and downregulated expression in *Nup210* KD cells. (e) Gene Ontology (GO) analysis of overlapped genes from (d). (f) qRT-PCR analysis of cell migration-related genes in *Nup210* KO and KD 4T1 cells. Multiple two tailed t-test, mean ± s.e.m, n=3 (sg-ctrl vs Nup210 KO), n=4 (sh-Ctrl vs sh-Nup210). (g) Representative ChIP-seq (NUP210-, H3K27Ac-, H3K27me3- and H3K4me3) tracks with predicted TADs within *Cxcl* and *Ccl2* region. (h) qRT-PCR analysis of cell migration-related genes in DMSO- and GSK126-treated *Nup210* KO 4T1 cells. Multiple two tailed t-test, mean ± s.e.m, n=3. (i) Representative images of H3.1/3.2 and H3K27me3 distribution in DMSO- and GSK126-treated *Nup210* KO cells. Scale bar = 5 μm. (j) H3.1/3.2 intensity ratio, H3K27me3 intensity and nuclear area quantification in DMSO- and GSK126-treated *Nup210* KO 4T1 cells. Kruskal-Wallis ANOVA with Dunn’s multiple comparison test, median with interquartile range. (k) Live cell tracking of nuclear area in DMSO- and GSK126 treated *Nup210* KO cells. N=40 (sg-Ctrl), n=41 (KO+DMSO), n=31.

Since enhancer-promoter interactions are critical for the regulation of gene expression^35, 36^ and H3.1/3.2 is preferentially enriched at the poised promoters, we applied structured illumination microscopy (SIM) to examine the intra-nuclear distribution of H3K4me3 (promoter) and H3K27Ac (enhancer) marks in *Nup210* KO cells. Although there was no significant changes in H3K4me3 volume in *Nup210* KO cells, H3K27Ac volume was significantly reduced (Extended Data Fig. 6e, f, Supplementary Video 3, 4) suggesting the decrease of enhancer regions. There was no noticeable alteration of H3K4me3 distribution in relation to H3.1/3.2 foci in *Nup210* KO cells (Extended Data Fig. 6g). However, ChIP-seq of H3K4me3 revealed that the TSS of 1199 genes showed 2-fold decreased enrichment of H3K4me3 in *Nup210* KO cells (Fig. 5c). RNA-seq of *Nup210* KD 4T1 cells identified 282 downregulated (2-fold) and 249 upregulated (2-fold) genes (Extended Data Fig. 6i), 62 of which had H3K4me3 loss at the promoter and ≥ 2-fold decreased expression (Fig. 5d) and thus were likely to be direct targets of NUP210 regulation. Gene Ontology (GO) analysis on the 62 genes showed enrichment of cell migration and chemotaxis processes (Fig. 5e), including chemokines (*Ccl2*, *Cxcl1*, *Cxcl3*), cytokine (*Il1a*), and cell adhesion (*Postn*, *Serpine1*, *Itga7*, *Itgb2*) molecules (Fig. 5f, Extended Data Fig. 6j). Similar transcriptional changes were also observed in NUP210-depleted 6DT1 cell line (Extended Data Fig. 6k). Integration of publicly available Hi-C data from mouse embryonic stem cells (mESC)^37^ using the 3D-Genome Browser^38^ revealed that H3K27me3 accumulated across the *Cxcl* and *Ccl2* loci topologically associated domains (TADs) in *Nup210* KO cells (Fig. 5g), and was accompanied by a marked loss of H3K4me3 at the promoters of these genes, implying that NUP210 loss results in heterochromatinization across the *Cxcl*/*Ccl2* enhancer and promoter regions.

To test whether the expression of NUP210-regulated genes can be rescued in *Nup210* KO cells by inhibiting heterochromatization, cells were treated with an EZH2 inhibitor GSK126 which prevents H3K27me3 modification. Treatment with GSK126 significantly increased the expression of *Ccl2*, *Cxcl1*, *Cxcl3*, *Postn*, and *Il-1a* (Fig. 5h), suggesting that they were suppressed through heteterochromatinization in *Nup210* KO cells. In contrast, GSK126 treatment decreased the expression of *Itga7*, *Itgb2*, and *Serpine1*, implying that they were either not directly suppressed by heteterochromatinization or could be suppressed by a non-specific effect of the drug. Interestingly, perinuclear distribution of H3.1/3.2 and nuclear size was significantly restored in GSK126-treated *Nup210* KO cells (Fig. 5i, j). Live-cell tracking of nuclear size further supported the observation that GSK126 treatment significantly rescued the nuclear size of *Nup210* KO cells (Fig. 5k). These results are consistent with the role of NUP210 in preventing H3K27me3 heterochromatinization of perinuclear DNA.

### Nup210 loss is associated with differential repositioning of cell migration-related gene loci

To determine whether NUP210 regulates its target genes through anchoring topologically associated domain (TADs) to the nuclear periphery, 3D-DNA FISH was performed. FISH probes were generated from BAC clones (∼2 Mb) spanning *Cxcl*, *Postn*, *Ccl2*, and *Itgb2* genomic regions. As expected, majority of the FISH spots were observed near the nuclear periphery in aneuploid 4T1 cells (Extended Data Fig. 7a). Interestingly, in *Nup210* KO cells, *Cxcl*, *Postn*, and *Itgb2* loci were repositioned within the nucleus (Extended Data Fig. 7b), with *Cxcl* and *Itgb2* loci repositioned closer to the nuclear periphery, and the *Postn* locus repositioned away from the periphery. Radial positioning of the *Ccl2* locus, however, was not significantly changed. Rather, the distance of *Ccl2* FISH spots to heterochromatin foci were significantly reduced in *Nup210* KO (Extended Data Fig. 7c) cells. These results suggest that NUP210 loss is associated with altered chromatin topology and nuclear positioning of cell migration-related gene loci. Since three dimensional chromatin domains are organized into transcriptionally active (A compartment) and repressive (B compartment) compartments within the nucleus^39^, repositioning of these genes in *Nup210* KO cells is consistent with their repositioning into repressive B compartment due to the heterochromatinization.

### NUP210 is mechanosensitive and regulates focal adhesion, cell migration, and invasion

Alterations in cell adhesion/migration-related gene expression can lead to changes in cellular adhesion, a critical phenotype in metastatic progression. *Nup210* KO cells exhibited less spread, round morphology when they were grown on type I collagen and fibronectin matrices (Fig. 6a). Staining for phospho-FAK (Y397) and phalloidin revealed that loss of *Nup210* significantly decreased the focal adhesions (area and count), actin stress fibers and cell spreading when plated on type I collagen or fibronectin (Fig. 6b, c). Similar results were obtained in case of *Nup210* knockdown 4T1 (Extended Data Fig. 8a), 6DT1 (Extended Data Fig. 8b, c) and human MDA-MB-231 cells (Extended Data Fig. 8d). Consistently, protein level of phospho-FAK (Y397) was also decreased in NUP210 depleted mouse (Extended Data Fig. 8e-g) and human (Extended Data. Fig. 8h, i) breast cancer cells. In addition, there was significant decrease of phospho-FAK (Y397) staining within the *Nup210* KO primary tumors suggesting that NUP210 loss also affects focal adhesion *in vivo* (Fig. 6d). To test whether the constitutively active FAK can rescue the metastasis defects of *Nup210* KO cells, we stably expressed Myc-tagged FAK in *Nup210* KO cells (Fig. 6e). Although there was no significant effect on primary tumors in FAK expressing cells (Fig. 6f), both tumor-normalized lung metastases count (Fig. 6h) and metastasis incidence (Fig. 6i) was significantly rescued indicating that NUP210 is mediating it’s effect on metastasis through focal adhesion signalling pathway.

**Figure 6:**
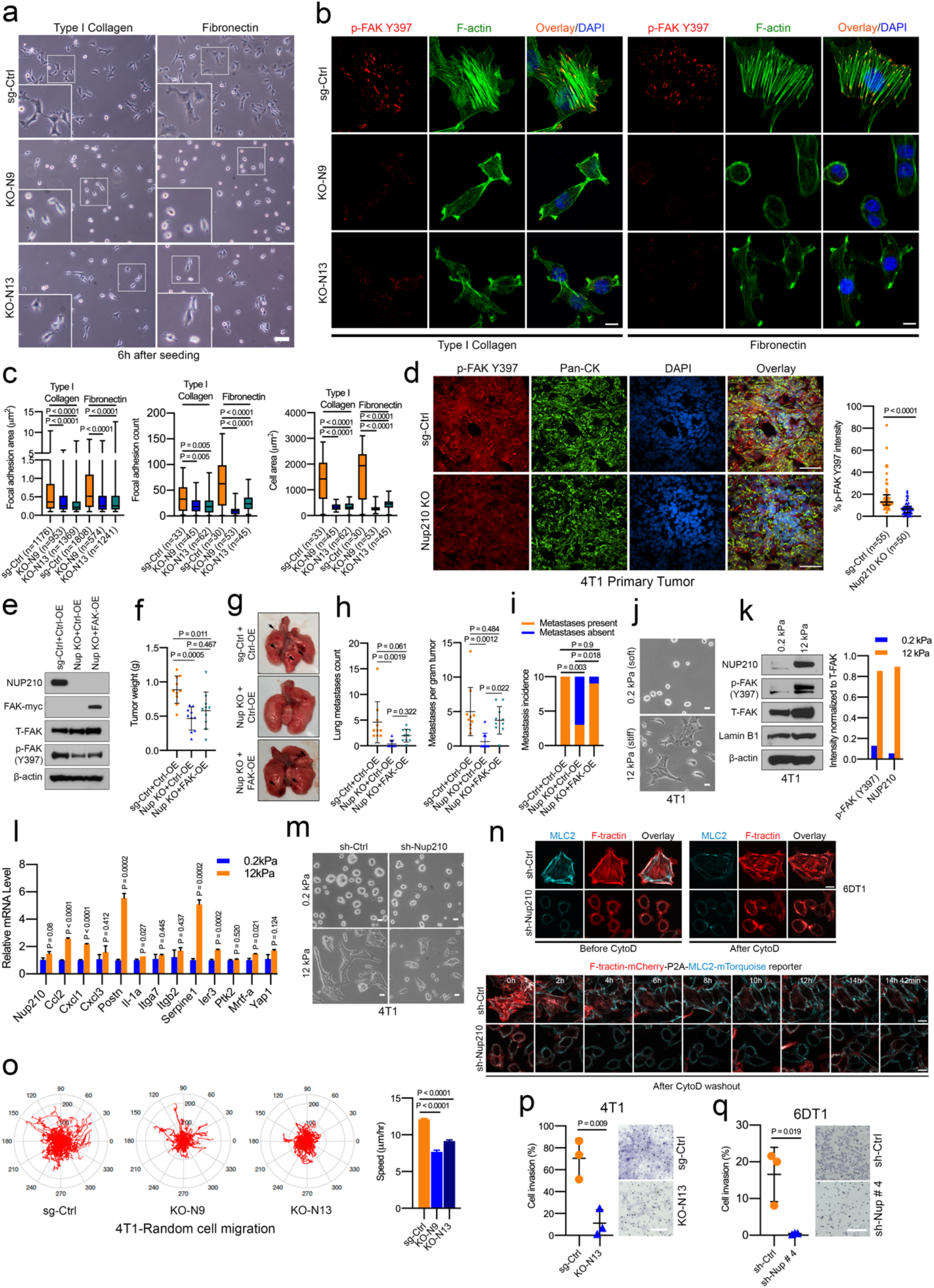
*Nup210* is mechanosensitive and regulates focal adhesion, cell migration, and invasion. (a) Representative brightfield images of 4T1 Nup210 KO cells grown on type I collagen and fibronectin. Scalebar = 100 μm. (b) Representative images of p-FAK (Y397) focal adhesion puncta and F-actin (Phalloidin) in sg-Ctrl and *Nup210* KO cells grown on type I collagen and fibronectin. Scale bar = 10 μm. (c) Quantification of focal adhesion area (left), focal adhesion count (middle) and cell spreading (right) in sg-Ctrl and *Nup210* KO 4T1 cells. ANOVA with Tukey’s multiple comparison test. For the boxplot, the box extends from 25^th^ to 75^th^ percentile, whiskers extend from smallest values to the largest values, horizontal line represents median. (d) (Left) Immunostaining of *Nup210* KO (KO-N13) 4T1 primary tumors with p-FAK Y397 antibody. Pan-Cytokeratin (Pan-CK) was used as tumor cell marker. Scalebar 50 μm. (Right) quantification of p-FAK Y397 signal in the Pan-CK-stained tumor area. N = 5 mice per condition, ∼10 independent field per mouse tumor section. Mann-Whitney U test, median with interquartile range. (e) Western blot of myc-tagged FAK overexpression in *Nup210* KO 4T1 cells. (f) Primary tumor weight in FAK overexpressing *Nup210* KO (KO-N13) 4T1 cells. ANOVA with Tukey’s multiple comparison test, mean ± s.d. (g) Representative lung images of the mice injected with FAK overexpressing *Nup210* KO cells. (h) (Left) Lung metastases count and (right) lung metastases normalized to tumor weight in FAK overexpressing *Nup210* KO cells. ANOVA with Tukey’s multiple comparison test, mean ± s.d. (i) Metastasis incidence in mice injected with FAK overexpressing *Nup210* KO cells. Chi-square test with Bonferroni correction. (j) Representative images of 4T1 cells on fibronectin-coated hydrogel layers of soft (0.2 kPa) and stiff (12 kPa) matrices. Scale bar = 10 μm. (k) (Left) Western blot of NUP210, p-FAK Y397, and T-FAK (Total FAK) proteins of 4T1 cells grown on soft or stiff matrices. (Right) Quantification of signals normalizd to T-FAK intensity. (l) qRT-PCR of *Nup210*-regulated genes as well as known mechanosensitive genes (*Ier3*, *Mrtf-a*, *Yap1*) in soft and stiff matrices. Multiple two tailed t-test, mean ± s.e.m. (m) Morphology of sh-Ctrl and *Nup210* KD 4T1 cells in soft and stiff matrices. Scale bar = 10 μm. (n) Live cell imaging of F-tractin-MLC2 reporter transduced 6DT1 *sh-Nup210* cells. Cells were treated with Cytochalasin D for 1hr and imaged overnight after drug washout. Scale bar = 10 μm. (o) (Left) cell migration tracks (red) of sg-Ctrl and *Nup210* KO 4T1 cells. (Right) bar graph of cell speed quantification. Kruskal-Wallis ANOVA with Dunn’s multiple comparison test, sg-Ctrl (n=847), KO-N9 (n=124), KO-N13 (n=375), mean ± s.e.m. (p) Quantification (left) and representative images (right) of cell invasion for 4T1 (at 48h) and (q) 6DT1 (at 24h) NUP210 depleted cells. Two tailed t-test, mean ± s.d. Scale bar = 100 μm.

As focal adhesion contributes to the mechanical response of tumor cells, we asked whether NUP210 is mechanosensitive in metastatic cells. 4T1 cells grown on soft/low stiffness (0.2 kPa) condition exhibited a round colony-like morphology, while on the higher stiffness conditions (12 kPa), these cells exhibited a more spread morphology (Fig. 6j). Increased protein levels of p-FAK (Y397) and NUP210 were found in the higher stiffness condition (Fig. 6k), indicating that NUP210 protein is mechanosensive in 4T1 cells. Moreover, NUP210 protein level was higher on cells grown on type I collagen than fibronectin (Extended Data Fig. 8j) suggesting that NUP210 might be sensitive to different ECM composition. Many of the NUP210-regulated genes were also sensitive to higher ECM stiffness condition (Fig. 6l). However, unlike protein level, transcript levels of Nup210, FAK (Ptk2) as well as known mechanosensitive transcription factor, Yap1 were not significantly upregulated in higher stiffness condition. This result is consistent with previous observation that soft matrices tend to degrade mechanosensitive proteins^40^ and rapid reduction of NUP210 protein on soft matrix could be due to the post-transcriptional loss of protein stability. NUP210 depleted cells exhibited decreased spreading on the higher stiffness condition (Fig. 6m), consistent with NUP210 contributing to mechanosensation. As mechanosensation can occur through alteration of actin dynamics^41, 42^, treatment with an actin polymerization inhibitor, Cytochalasin D in 4T1 cells demonstrated that NUP210 depleted cells had decreased recovery of actin polymerization, as marked by phalloidin staining of actin stress fibers (Extended Data Fig. 8k). As mechanical response is mediated through alteration of actomyosin tension, staining with phospho-myosin light chain 2 (MLC2-S19) and phalloidin revealed a decreased association of F-actin and activated myosin II (phospho-MLC2-S19) in *Nup210* KO cells grown on type I collagen and fibronectin matrices (Extended Data Fig. 8l). To further investigate the actomyosin contractility of *Nup210*-depleted cells, we stably expressed a fluorescently tagged stochiometric F-actin (Ftractin) and myosin II activity reporter (MLC2)^43^ in 6DT1 cells and performed live-cell imaging analysis. On type I collagen, both F-tractin and MLC2 remained associated at the actin stress fibers and lamellipodia in control cells compared to *Nup210* knockdown cells where they were localized at the nuclear periphery (Fig. 6n). In Cytochalasin D treated condition, Ftractin and MLC2 association was disrupted. After Cytochalasin D washout, control cells quickly recovered and maintained their Ftractin-MLC2 association at the lamellipodia (Supplementary Video 5). In contrast, this association was disrupted in *Nup210* knockdown cells and MLC2 mainly localized at the nuclear periphery (Supplementary Video 6). These results suggests that the loss of NUP210 is associated with decreased actomyosin contractility of cancer cells.

As actomyosin contractility is critical for cell migration and invasion, live-cell imaging analysis of cell migration was performed and revealed a significant decrease of cell migration in *Nup210* KO cells than sg-Ctrl cells (Fig. 6o, Supplementary Videos 7, 8, 9). A significant reduction of cell invasion was also observed in NUP210 depleted 4T1 (Fig. 6p) and 6DT1 cells (Fig. 6q). Taken together, these results demonstrate that NUP210 is part of a sensor of the extracellular matrix stiffness and composition that significantly affects the migratory and invasive ability of tumor cells.

### NUP210 is a LINC (Linker of nucleoskeleton and cytoskeleton) complex associated protein and regulates mechanosensitive gene expression

As mechanotransduction is mediated through the nuclear envelope-anchored LINC complex proteins^42, 44^, we asked whether NUP210-dependent mechanical response is mediating through its association with LINC complex proteins. Co-IP revealed that NUP210 preferentially interacts with LINC complex protein SUN2, but not SUN1 (Fig. 7a). Immunofluorescence imaging showed NUP210 was associated with SUN2 at the nuclear periphery (Fig. 7b). Interestingly, perinuclear localization of SUN1 and SUN2 were significantly disrupted in *Nup210* KO cells (Fig 7c). SUN2 was distributed diffusedly throughout the cells with significant increase in intranuclear distribution in *Nup210* KO cells (Fig. 7d). In addition, the ratio of nuclear periphery to total nuclear intensity was significantly decreased in case of both SUN2 and SUN1. Since NUP210 was affecting LINC complex, we looked at the nuclear lamina of *Nup210* KO cells more closely. Although there was no significant changes in total Lamin B1 intensity, total Lamin A/C intensity was significantly increased in *Nup210* KO cells (Fig. 7e, f). In contrast, the ratio of nuclear periphery to total intensity of both Lamin B1 and Lamin A/C was significantly reduced in *Nup210* KO cells. More interestingly, 3D surface reconstruction revealed that both Lamin B1 and Lamin A/C appeared as crevices inside the nucleus of *Nup210* KO cells (Fig. 7g, Supplementary Video 10, 11). Since increased Lamin A/C level has previously been shown to increase nuclear stiffness resulting in decreased cell migration and tumorigenesis^45, 46^, we measured the level of total Lamin A/C protein using western blot. However, total Lamin A/C protein level was not significantly changed in *Nup210* KO cells (Extended Data Fig. 9a) suggesting that the increased Lamin A/C intensity in immunofluorescence was probably due to the heterochromatin-mediated decrease of nuclear size which resulted in higher concentration of Lamin A/C at the nuclear envelope. Subcellular fractionation revealed that SUN2 and Lamin B1 proteins are mislocalized in the cytoplasmic fraction of *Nup210* KO cells (Fig. 7h). There was overall decrease of SUN1 protein in the nuclear fraction of *Nup210* KO cells. Nuclear translocation of known mechanosensitive proteins (MRTF-A and YAP) were not affected in *Nup210* KO cells (Fig. 7h). Rather, there was overall decrease of MRTF-A/MKL1 in nuclear fraction and level of YAP was decreased in both cytoplasmic and nuclear fraction of *Nup210* KO cells. This result suggests that the role of NUP210 in mechanosensation is independent of the canonical nuclear translocation mechanism of mechanosensitive proteins^41, 47^. However, overall distribution of nuclear pore was not affected by *Nup210* KO (Extended Data Fig. 9b). In addition, dynamic organization of nuclear pore was not compromized in NUP210 depleted cells as confirmed by FRAP analysis of GFP-tagged nuclear pore protein POM121 (Extended Data Fig. 9c). These results suggested that the loss of NUP210 is associated with alteration of nuclear lamina and uncoupling of LINC complex machinery resulting in altered mechanical response, cell migration and metastasis.

**Figure 7:**
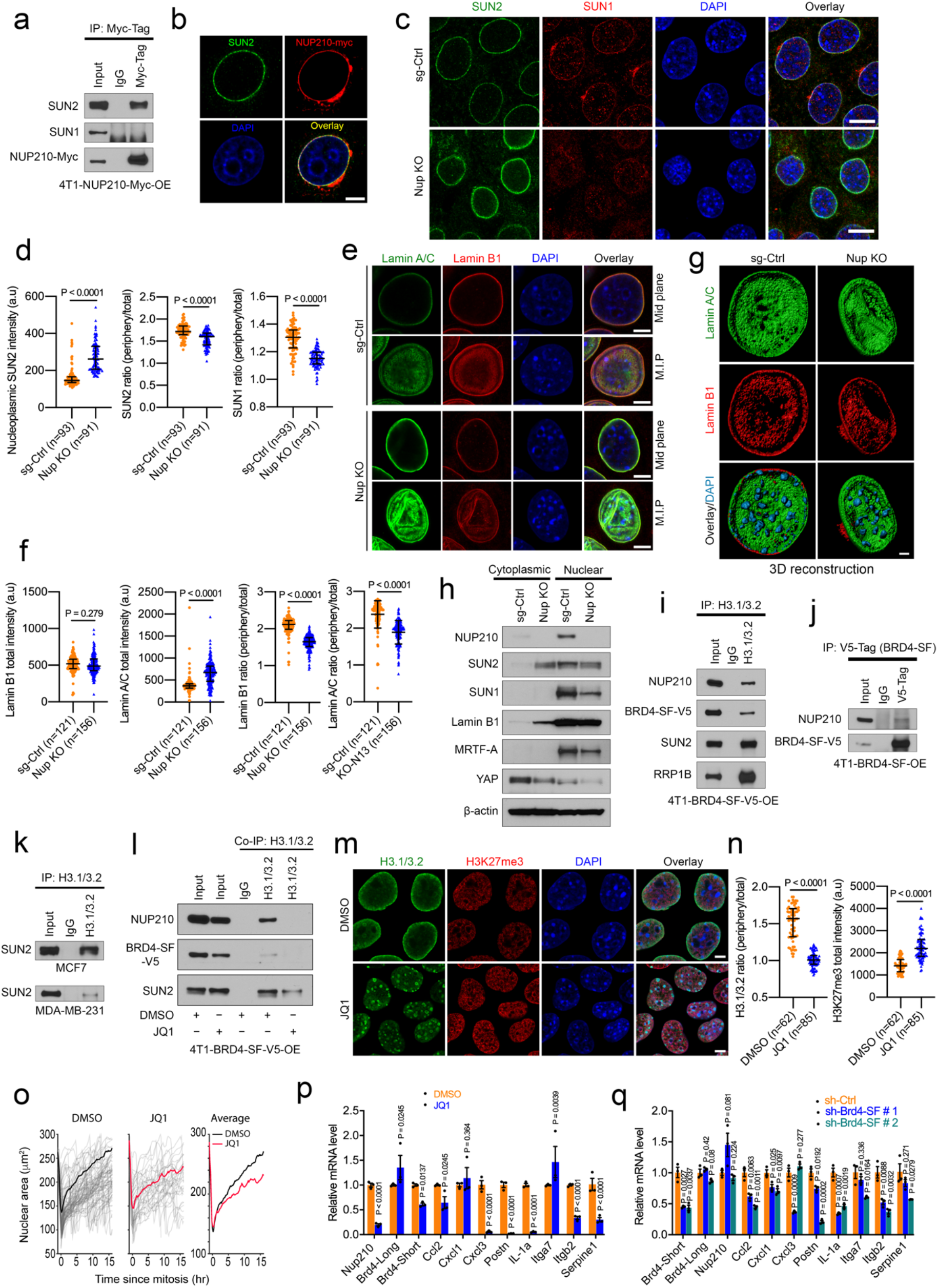
NUP210 is a LINC complex associated protein that mediates chromatin connection through the short isoform of BRD4. (a) Coimmunoprecipitation of myc-tagged NUP210 with LINC complex protein SUN1 and SUN2 (b) Colocalization of SUN2 and myc-tagged NUP210 at the nuclear periphery. Scale bar = 5 μm. (c) Distribution of SUN1 and SUN2 in *Nup210* KO (KO-N13) 4T1 cells. Scale bar = 10 μm. (d) Quantification of SUN2 and SUN1 intensity in *Nup210* KO 4T1 cells. Mann-Whitney U test, median with interquartile range. (e) Distribution of Lamin B1 and Lamin A/C in *Nup210* KO 4T1 cells. M.I.P = Maximum intensity projection. Scale bar = 5 μm. (f) 3D reconstruction of Lamin B1, Lamin A/C and DAPI (heterochromatin foci) distribution in *Nup210* KO 4T1 cells. Scale bar = 2 μm. (g) Quantification of Lamin B1 and Lamin A/C intensity in *Nup210* KO 4T1 cells. Mann-WhitneyU test, median with interquartile range. (h) Subcellular fractionation of LINC complex proteins and known mechanosensitive proteins (MRTF-A and YAP) in *Nup210* KO 4T1 cells. (i) Co-IP showing H3.1/3.2 interactions with LINC complex proteins SUN2, BRD4 short isoform (BRD4-SF) and RRP1B in 4T1 cells. (j) Co-IP showing the interaction of V5-tagged BRD4-SF with NUP210 in 4T1 cells. (k) Co-IP showing H3.1/3.2 interaction with SUN2 in human MCF7 and MDA-MB-231 cell line. (l) Co-IP showing the disruption of H3.1/3.2 interaction with NUP210 and SUN2 in JQ1-treated 4T1 cells. (m) Distribution of H3.1/3.2 and H3K27me3 in 4T1 cells treated with bromodomain inhibitor JQ1. Scale bar 5 μm. (n) Quantification of H3.1/3.2 and H3K27me3 intensity in JQ1-treated 4T1 cells. Mann-Whitney U test, median with interquartile range. (o) Live-cell tracking of nuclear size in JQ1-treated 4T1 cells. For DMSO, n = 122 and for JQ-treated condition, n = 127. (p) qRT-PCR of NUP210-regulated genes in JQ1-treated 4T1 cells. Multiple two tailed t-test, mean ± s.e.m. (q) qRT-PCR of NUP210-regulated genes in BRD4-SF knockdown 4T1 cells. Multiple two tailed t-test, mean ± s.e.m.

We then investigated the link between the alteration of mechanotransduction machinery and transcriptional suppression of *Nup210* KO cells. We have previously shown that metastasis susceptibility genes RRP1B and the short isoform of BRD4 (BRD4-SF) interacts with LINC complex proteins at the nuclear lamina^19, 48^. Similarly, BRD4 has been linked to mechanotransduction^49^ and metastasis^50^. So, we hypothesized that NUP210 is probably associated with this protein complex and mediates the interaction with H3.1/3.2-associated chromatin to regulate the gene expression in metastasis. Co-IP revealed that endogenous histone H3.1/3.2 was associated with NUP210, BRD4-SF, SUN2 and RRP1B protein complex in 4T1 cells (Fig. 7i). BRD4-SF also interacts with NUP210 in these cells (Fig. 7j). H3.1/3.2-SUN2 interaction was also evident in human breast cancer cells MCF7 and MDA-MB-231 (Fig. 7k). We then tested the necessity of either NUP210 or BRD4-SF in bridging the interaction between the LINC complex machinery and H3.1/3.2 on chromatin. Although the interaction of SUN2 with H3.1/3.2 was unperturbed in *Nup210* KO cells (Extended Data Fig. 10a), complete disruption of H3.1/3.2-NUP210 interaction and partial disruption of H3.1/3.2-SUN2 interactions were observed in bromodomain inhibitor JQ1 treated 4T1 cells (Fig. 7l). This result suggested that BRD4-SF is crucial for bridging the interactions among the molecules in this putative complex. Interestingly, similar to *Nup210* KO, redistribution of H3.1/3.2 from the nuclear periphery to the interior and increased heterochromatin mark H3K27me3 was observed in bromodomain inhibitor JQ1 treated cells (Fig. 7m, n). Live-cell imaging of nuclear size also revealed that JQ1 treatment significantly decreased nuclear size (Fig. 7o). Decreased expression of *Nup210* and some of the *Nup210*-dependent genes was found in JQ1 treated cells (Fig. 7p). Similar gene expression changes were also observed in BRD4-SF specific knockdown 4T1 cells (Fig. 7q). Taken together, NUP210 is a LINC complex associated protein and suggests NUP210 is a regulator of mechanosensitive gene expression program through interacting with BRD4-SF-H3.1/3.2-associated chromatin.

To determine the cause and effect relationship of NUP210 dependent response, we treated 4T1 cells with actin polymerization inhibitor cytochalasin D (CytoD) and a global heterochromatin promoting H3K27me3-specific histone demethylase inhibitor (GSKJ4). Actomyosin organization was drastically reduced in CytoD treated cells as shown by F-actin and p-MLC2-S19 immunostaining (Extended Data Fig. 10b). Unlike CytoD, GSKJ4 had a minor effect on F-actin stress fibers but it disrupted p-MLC2-S19 distribution suggesting the alteration of actomyosin tension in GSKJ4-treated cells. Treatment with both of these drugs resulted in redistribution of histone H3.1/3.2 from the nuclear periphery to heterochromatin foci, similar to the effect seen in *Nup210* KO cells (Extended Data Fig. 10c, d). Furthermore, CytoD treatment showed additional effects such as micronuclei and double nuclei formation. Treatment of cells with GSKJ4 resulted in a similar transcriptional profile as *Nup210* depleted cells, which was not recapitulated by the treatment with CytoD (Extended Data Fig. 10e). *Nup210* mRNA level was slightly decreased in CytoD treated cells but unchanged at the protein level (Extended Data Fig. 10f). Despite the unchanged level of *Nup210* mRNA in GSKJ4-treated cells, a significant decrease of NUP210 protein level was observed suggesting post-transcriptional downregulation of *Nup210* (Extended Data Fig. 10f). Moreover, decreased levels of focal adhesion (p-FAK, T-FAK) and LINC complex protein (SUN2) were found in the GSKJ4-treated condition. Decreased Lamin B1 protein level by both drugs indicated that they are also capable of disrupting the nuclear lamina. Taken together, these results suggest a positive feedback loop between NUP210-dependent heterochromatin regulation and actin cytoskeletal tension, and the change in focal adhesion is likely to be the consequence of disrupting this feedback loop (Extended Data Fig. 10g).

### NUP210 regulates circulating tumor cells in mice

Many of the NUP210-regulated genes are secretory molecules like cytokines (IL-1*α*), chemokines (CCL2, CXCL1, CXCL3), and adhesion molecules (POSTN, SERPINE1), some of which have known pro-metastatic roles in breast cancer^51–54^. Cytokine array analysis revealed decreased secretion of cytokines (IL-1α), chemokines (CCL2, CXCL1), and adhesion molecules (POSTN, SERPINE1) in *Nup210* knockdown 4T1 cells (Fig. 8a). To investigate whether NUP210 mediates its effect on focal adhesion and cell migration through transcriptional control of secretory molecules, *Ccl2* was chosen as a candidate and examined for NUP210 KO-associated phenotypes. Knockdown of *Ccl2* in 4T1 cells (Fig. 8b) resulted in decreased p-FAK (Y397) (Fig. 8c), decreased focal adhesion and cell spreading (Fig. 8d, e). Moreover, a significant decrease in cell migration was observed upon *Ccl2* knockdown phenocopying the loss of NUP210 in these cells (Fig. 8f).

**Figure 8:**
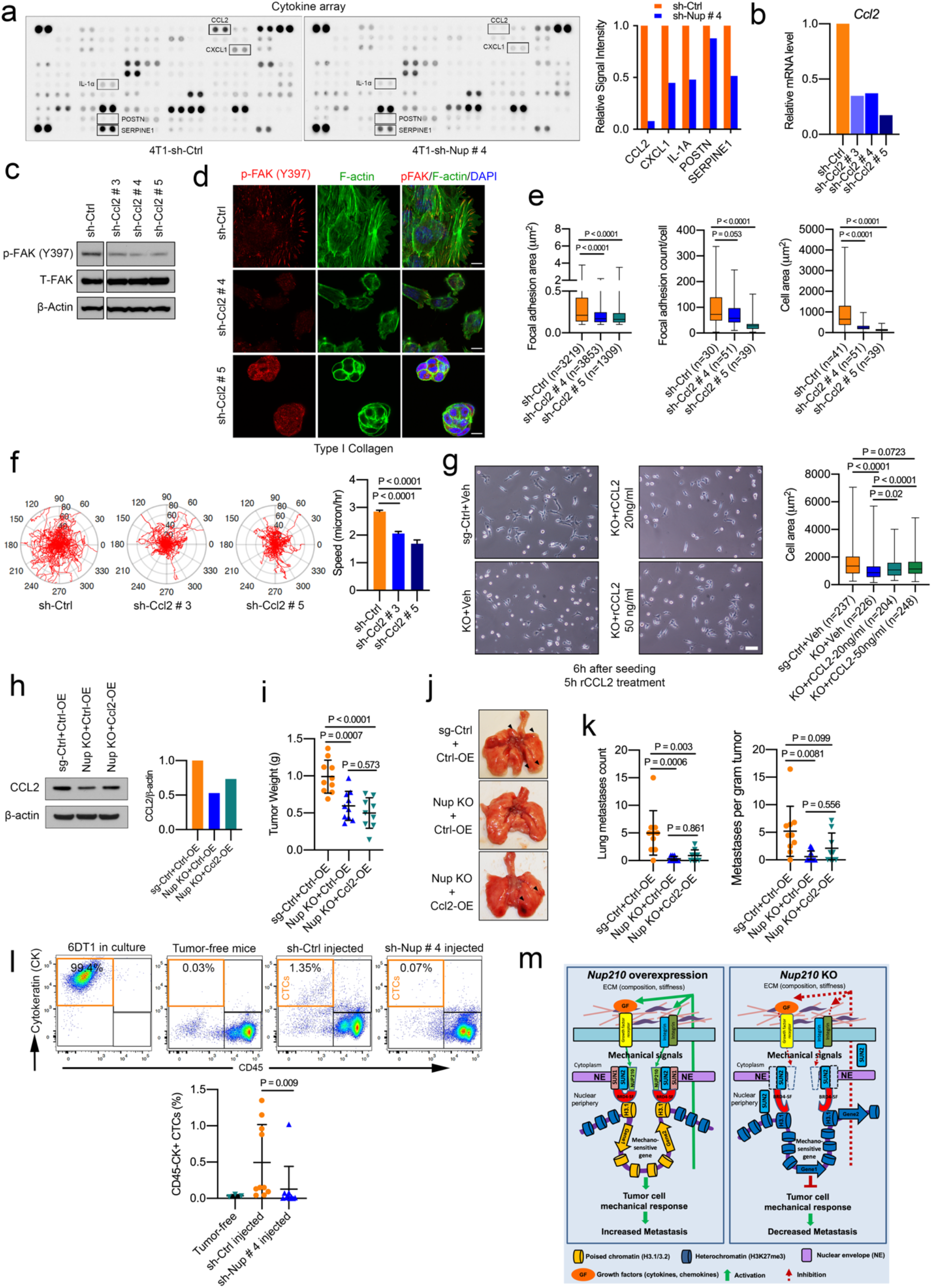
NUP210 regulates circulating tumor cells in mice. (a) (Left) Cytokine array on culture supernatants from sh-Ctrl and *Nup210* KD 4T1 cells. (Right) Quantification of the signals of each cytokines/chemokines. (b) qRT-PCR analysis of *Ccl2* KD in 4T1 cells. (c) Western blot of p-FAK Y397 and T-FAK proteins in *Ccl2* KD 4T1 cells. (d) p-FAK (Y397) focal adhesion puncta and F-actin distribution in sh-Ctrl and *Ccl2* KD 4T1 cells. Scale bar = 10 μm. (e) Quantification of focal adhesion area (left), adhesion number (middle) and cell spreading (right) in sh-Ctrl and *Ccl2* KD 4T1 cells. ANOVA with Tukey’s multiple comparison test, the box extends from 25^th^ to 75^th^ percentile, whiskers extend from smallest values to the largest values, horizontal line represents median. (f) (Left) Cell migration tracks (red) and quantification (right) of sh-Ctrl and *Ccl2* KD 4T1 cells. ANOVA with Tukey’s multiple comparison test, mean ± s.e.m. (g) (Left) Brightfield images showing recombinant Ccl2 treatment effect on cell spreading of *Nup210* KO 4T1 cells and (Right) quantification of cell area. ANOVA with Tukey’s multiple comparison test, the box extends from 25^th^ to 75^th^ percentile, whiskers extend from smallest values to the largest values, horizontal line represents median. (h) (Left) Western blot of *Ccl2* overexpression in *Nup210* KO 4T1 cells and (right) quantification of signal. (i) Tumor weight of mice injected with *Ccl2* overexpressing *Nup210* KO 4T1 cells. ANOVA with Tukey’s multiple comparison test, mean ± s.d. (j) Representative lung images from the mice mentioned in (i). (k) (Left) Lung metastases count and (right) lung metastasis normalized to tumor weight from the mice injected with *Ccl2* overexpressing *Nup210* KO 4T1 cells. ANOVA with Tukey’s multiple comparison test, mean ± s.d. (l) (Top) Flow cytometry analysis of the percentage of CD45-/CK+ circulating tumor cells (CTCs) and (bottom) quantification of CTCs in *Nup210* KD 6DT1 cell-injected mice blood. (m) Proposed model of NUP210 function in metastasis.

We then tested whether recombinant CCL2 can rescue the spreading defect of *Nup210* KO cells. Treating with recombinant CCL2 for 5 hours moderately rescued the spreading phenotype of *Nup210* KO cells (Fig. 8g). To test whether CCL2 can rescue the metastatic defect of *Nup210* KO cells, we overexpressed *Ccl2* in *Nup210* KO cells through lentiviral system. However, only ∼25% of *Ccl2* expression was rescued (Fig. 8h). Injecting these *Ccl2* overexpressing cells into mice did not have significant effect on primary tumors (Fig. 8i). Although *Ccl2* expression in *Nup210* KO cells showed increased trend of lung metastasis, it was not statistically significant (Fig. 8j, k). This result suggests that either the level of rescue of *Ccl2* expression in *Nup210* KO cell was not sufficient to have a significant effect on metastasis or rescuing with a single gene from the NUP210 transcriptional network cannot fully recover the NUP210 loss.

As *Nup210*-depleted cancer cells exhibited adhesion defects and decreased the level of secretory molecules necessary for tumor cell adhesion to endothelial wall^55^, we speculated that NUP210 loss might affect the circulating tumor cell (CTC) levels. Therefore, based on CD45-/CK+ staining, putative CTCs were isolated 28 days after orthotopic implantation of 6DT1 cells^56^. Under these conditions, approximately ∼0.5-1% of the cells in the blood of 6DT1-sh-Ctrl tumor-bearing mice were CD45-/CK+ putative CTCs. In contrast, CTCs counts in mice with *Nup210*-depleted tumors were indistinguishable from the FVB tumor-free control (Fig. 8l). These results suggest that the effect on focal adhesion and cell migration in NUP210 depleted cells is mediated through the NUP210-dependent transcriptional control of cytokine/chemokine secretory pathways that are required for tumor cell extravasation or for the survival of cells within the bloodstream.

In summary, our model suggests that growth factors and integrins transmit the mechanical signals via NUP210-SUN1-SUN2-BRD4-SF-H3.1/3.2 protein complex to regulate mechanosensitive genes (Fig. 8m), which in turn activate downstream signaling pathways necessary for focal adhesion, cell migration and metastasis. In *Nup210* KO cells, this signaling cascase is disrupted, thereby decreasing metastasis.

## DISCUSSION

Apart from providing structural support and nucleocytoplasmic transport^57^, the role of the nuclear pore complex in developmental gene regulation has drawn much attention recently^58–61^. Although NUP210 has been implicated in muscle differentiation^28, 62^, cellular reprogramming^63^, and T cell signaling^64, 65^, its function in human cancer was unknwon. Here, we have identified *Nup210* as a potential metastasis susceptibility gene for human ER+ breast cancer patients and demonstrated a previously unrecognized role of nuclear pore proteins in sensing the mechanical stress of the extracellular matrix.

The present study has important implications in the field of cellular mechanosensing. The nucleus is regarded as a sensor of mechanical stress where nuclear envelope-anchored, LINC complex proteins are thought to serve as mechanotransducer to regulate gene expression^44^. Despite being embedded in the nuclear envelope, direct involvement of nuclear pore proteins in mechanosensation remains unclear. Previous studies have focused mainly on the translocation of mechanosensitive transcription factors to the nucleus in a nucleocytoplasmic transport-dependent manner^40, 41, 47^. In contrast, our study has identified a transport independent role of nuclear pore components in regulating mechanosensitive, pro-metastatic genes through interaction with LINC complex proteins and chromatin. However, we could not specifically rule out the causal effect of NUP210 loss due to the potential positive feedback loop between heterochromatin regulation and actin cytoskeletal tension. Further studies will be necessary to fully understand the mechanism of NUP210 in regulating cellular mechanosensation.

Intriguingly, many of the NUP210-regulated genes were immune molecules like chemokines and cytokines. This result has important implications since rapid transcriptional activation and secretion of immune molecules are required to induce an immune response^66^. Consistent with T-cell defects in NUP210 deficient mice^64^, the nuclear pore complex might be essential for inducing an immune response. During metastasis, cancer cells may exploit this mechanism for robust transcription of these genes, which then promote their migration into distant organs in an autocrine fashion.

Mechanistically, we have demonstrated that the loss of NUP210 is associated with decreased peripheral localization of H3.1/3.2 and increased enrichment near heterochromatin foci. In support of this observation, we found increased accumulation of H3K27me3-marked heterochromatin in *Nup210* KO cells. These results suggest a role for NUP210 as a chromatin barrier insulator-binding protein. The nuclear pore complex has been described as a heterochromatin barrier component of the nuclear lamina^67, 68^. Loss of barrier insulator elements is associated with increased spreading of heterochromatin marks to adjacent actively transcribing gene loci^69^. Since *Nup210* loss was associated with H3K27me3-marked heterochromatin spreading, NUP210 is probably acting as a barrier of H3K27me3-mediated heterochromatin spread to adjacent loci at the nuclear pore.

The potential enrichment of NUP210 at active enhancers and its association with chromatin reader BRD4-SF is consistent with the previous observation that the oncogenic properties of BRD4-SF is partly mediated through enhancer regulation^50^. Nuclear pores can be a scaffold of enhancer-promoter contact mediating transcriptional activation of poised genes^61^. It is therefore conceivable that in response to higher stiffness of the extracellular microenvironment, NUP210 can facilitate the interaction of LINC complex with BRD4-SF and histone H3.1/3.2-marked enhancer regions to establish distal regulatory element interaction with poised promoters of mechanosensitive genes necessary for metastatic progression.

Treating *Nup210* KO cells with an EZH2 inhibitor (GSK126) could partially rescue the peripheral distribution of H3.1/3.2 and mechanosensitive gene expression. This has important clinical implications since EZH2 is a therapeutic target for multiple malignancies and some EZH2 inhibitors are currently in clinical trials^70, 71^. In breast cancer, both tumor-promoting and inhibitory effects have been described for EZH2 in a context-specific manner^72^. Treating with an EZH2 inhibitor could adversely affect patient outcome through activation of the mechanosensitive genes. In line with our observation, GSK126 treatment has been shown to exhibit adverse effects via inhibiting anti-tumor immunity^73^ and promoting inflammation^74^ in several pre-clinical models of cancer.

In conclusion, we have identified a previously unrecognized role for nuclear pore protein, NUP210 in mechanical sensing of aggressive metastatic cancer cells. Future studies should determine whether blocking NUP210-chromatin interactions can be a therapeutic strategy to prevent metastasis.

## ACKNOWLEDGEMENTS

The authors thank Sarah Deasy, Ngoc-Han Ha and Brandi Carofino for critical review of the manuscript, Center for Cancer Research (CCR) Flow Cytometry Facility member Ferenc Livak for FACS analysis, CCR sequencing facility (Frederick, MD) for RNA-seq/ChIP-seq analysis, CCR mass spectrometry core facility for LC-MS analysis. The study was supported by the Intramural Research Program, National Cancer Institute, NIH.

## AUTHOR CONTRIBUTIONS

R.A conceived the project, performed the majority of the experiments, analyzed the data, and wrote the manuscript. A.S, J.J.Z, S.K, P.W, and S.Z.T performed chromatin accessibility, ChIA-PET analysis, and wrote the manuscript. S.B performed DNA FISH analysis. H.L and M.P.L performed ChIP-seq/RNA-seq analysis. D.P and S.D.C analyzed live-cell imaging data. A.T and M.J.K performed microscopy image analysis and wrote the manuscript. J.E.D and R.M.S analyzed immunohistochemistry data. G.L.H supervised the chromatin accessibility analysis. Y.R supervised the ChIA-PET analysis. K.W.H conceived and supervised the project, analyzed the data, and wrote the manuscript.

## EXTENDED DATA FIGURE LEGENDS

**Extended Data Figure 1:**
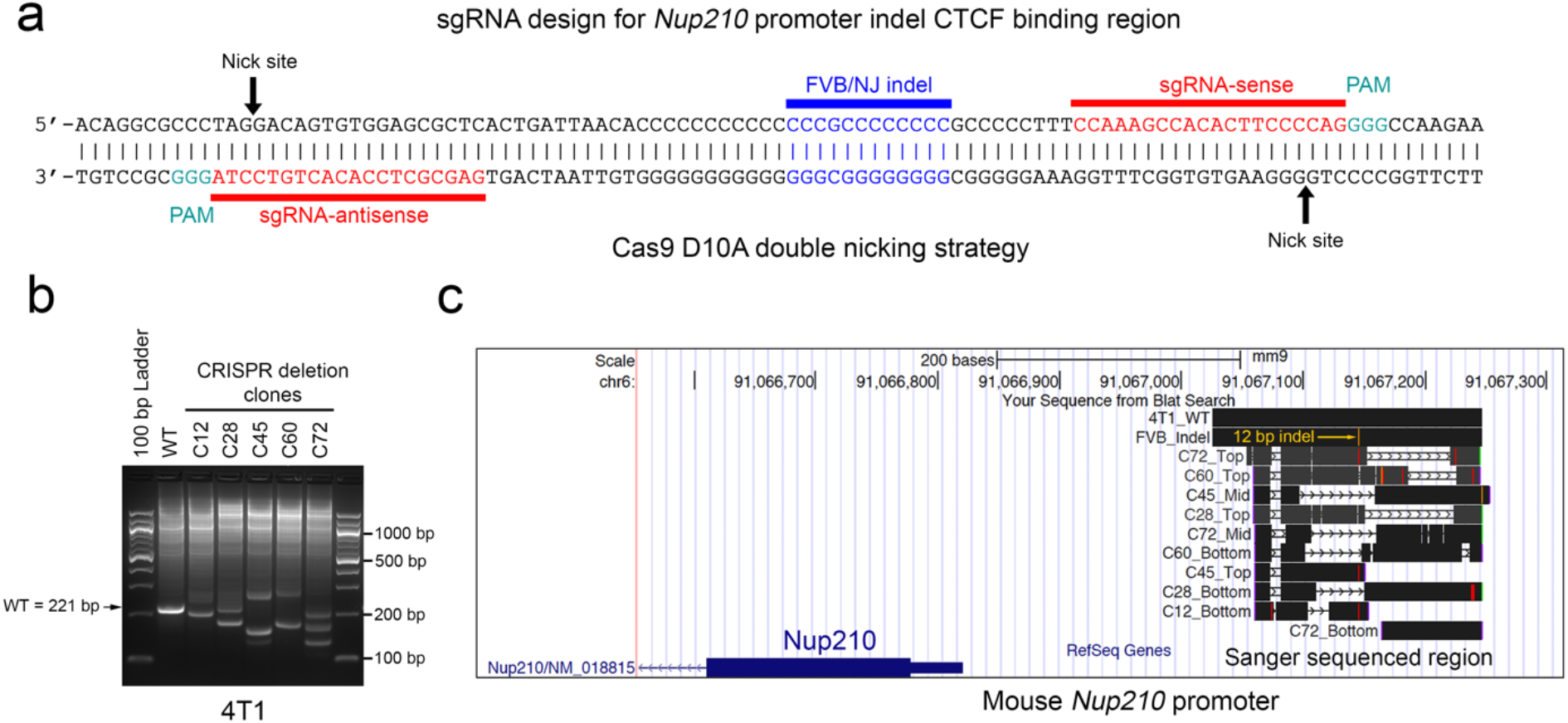
CRISPR/Cas9-mediated deletion of CTCF binding site on *Nup210* promoter. (a) sgRNA design strategy for the deletion of putative CTCF binding site on Nup210 promoter of 4T1 (BALB/cJ-derived) cells. FVB/NJ 12 bp indel region on CTCF binding site was shown in blue. (b) PCR amplification of 221 bp region spanning the Cas9 D10A-deleted CTCF binding site in 4T1 cell clones. (c) UCSC genome browser view of the DNA sequences of individual band (top/mid/bottom) obtained from each clone mentioned in (b). Line with arrow indicates mutated region.

**Extended Data Figure 2:**
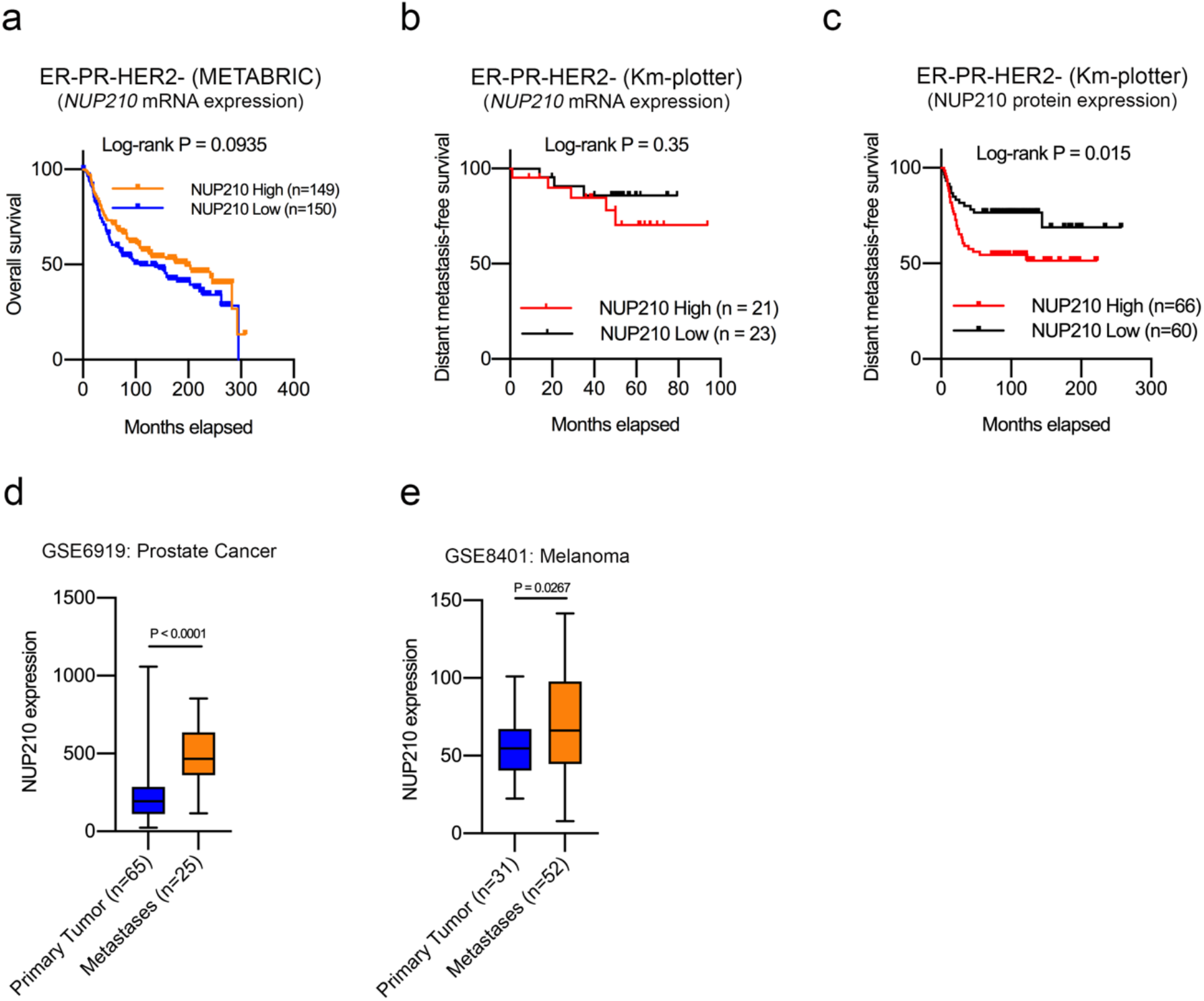
Association of NUP210 expression in triple negative (ER-/PR-/HER2-) patient and NUP210 expression in human metastases datasets. (a) Association of NUP210 mRNA with overall survival and (b) DMFS in ER-/PR-/HER2-patients. (c) Association of NUP210 protein level on distant metastasis-free survival of ER-/PR-/HER2-patient. (d) Human prostate cancer dataset showing the differential expression of *NUP210* mRNA between primary tumor and metastases. Mann-Whitney U test, the box extends from 25^th^ to 75^th^ percentile, whiskers extend from smallest values to the largest values, horizontal line represents median. (e) Human melanoma dataset showing the differential expression of *NUP210* mRNA between primary tumor and metastases. Mann-Whitney U test, the box extends from 25^th^ to 75^th^ percentile, whiskers extend from smallest values to the largest values, horizontal line represents median.

**Extended Data Figure 3:**
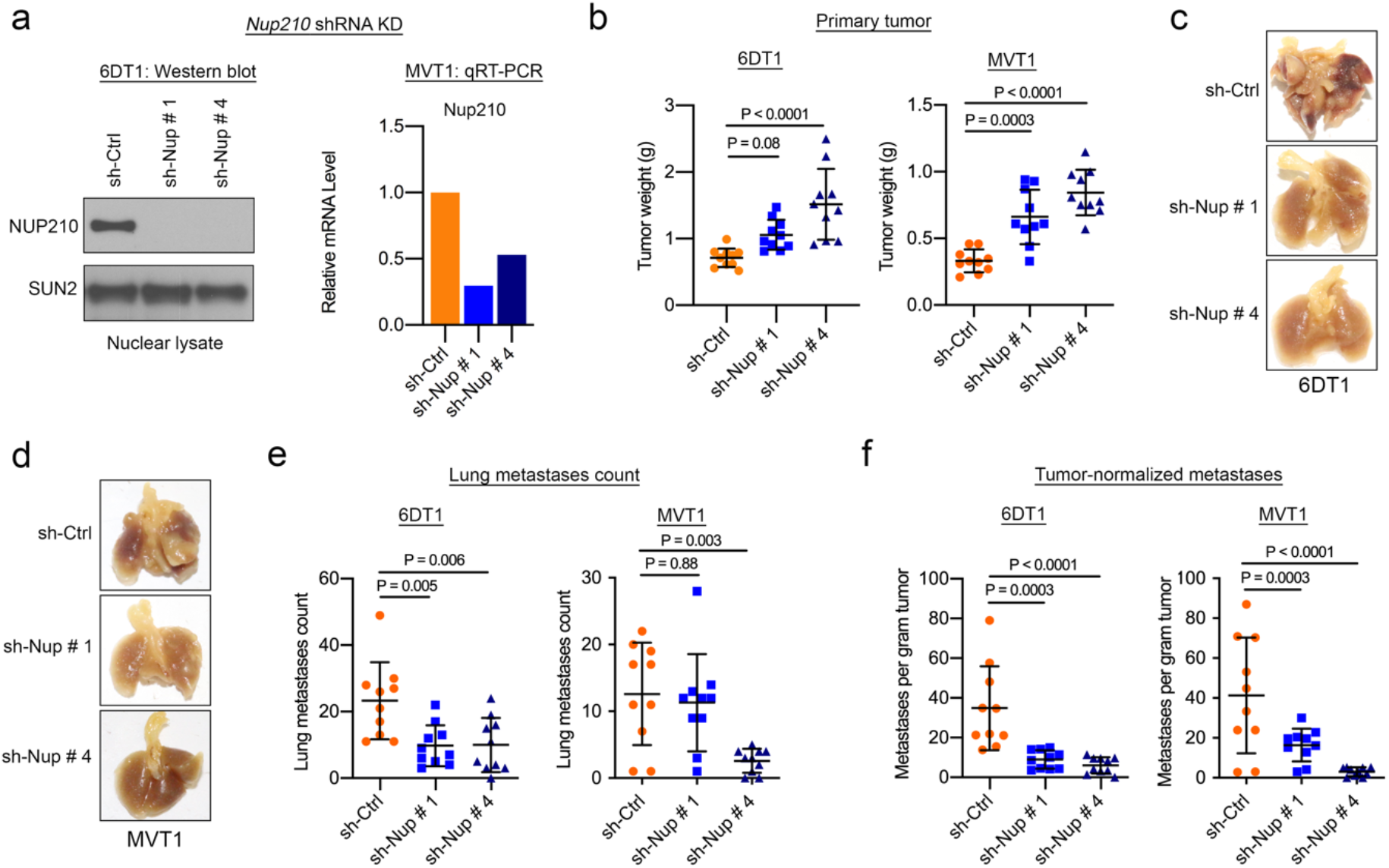
Knockdown of *Nup210* in the 6DT1 and MVT1 cell lines decreases lung metastasis. (a) (Left) western blot NUP210 protein of *Nup210* KD 6DT1 cells and (right) qRT-PCR of *Nup210* KD level in MVT1 cells. (b) Primary tumor weight after orthotopic transplantation of *Nup210* KD 6DT1 (left) or MVT1 (right) cells. ANOVA with Tukey’s multiple comparison test, mean ± s.d. (c) Representative lung images of the mice injected with *Nup210* KD 6DT1 cells. (d) Representative lung images of the mice injected with *Nup210* KD MVT1 cells. (e) Lung metastases count after orthotopic transplantation of *Nup210* knockdown 6DT1 (left) and MVT1 (right) cells. ANOVA with Tukey’s multiple comparison test, mean ± s.d. (f) Lung metastases count normalized to primary tumor weight, derived from values shown in (b) and (e). ANOVA with Tukey’s multiple comparison test, mean ± s.d.

**Extended Data Figure 4:**
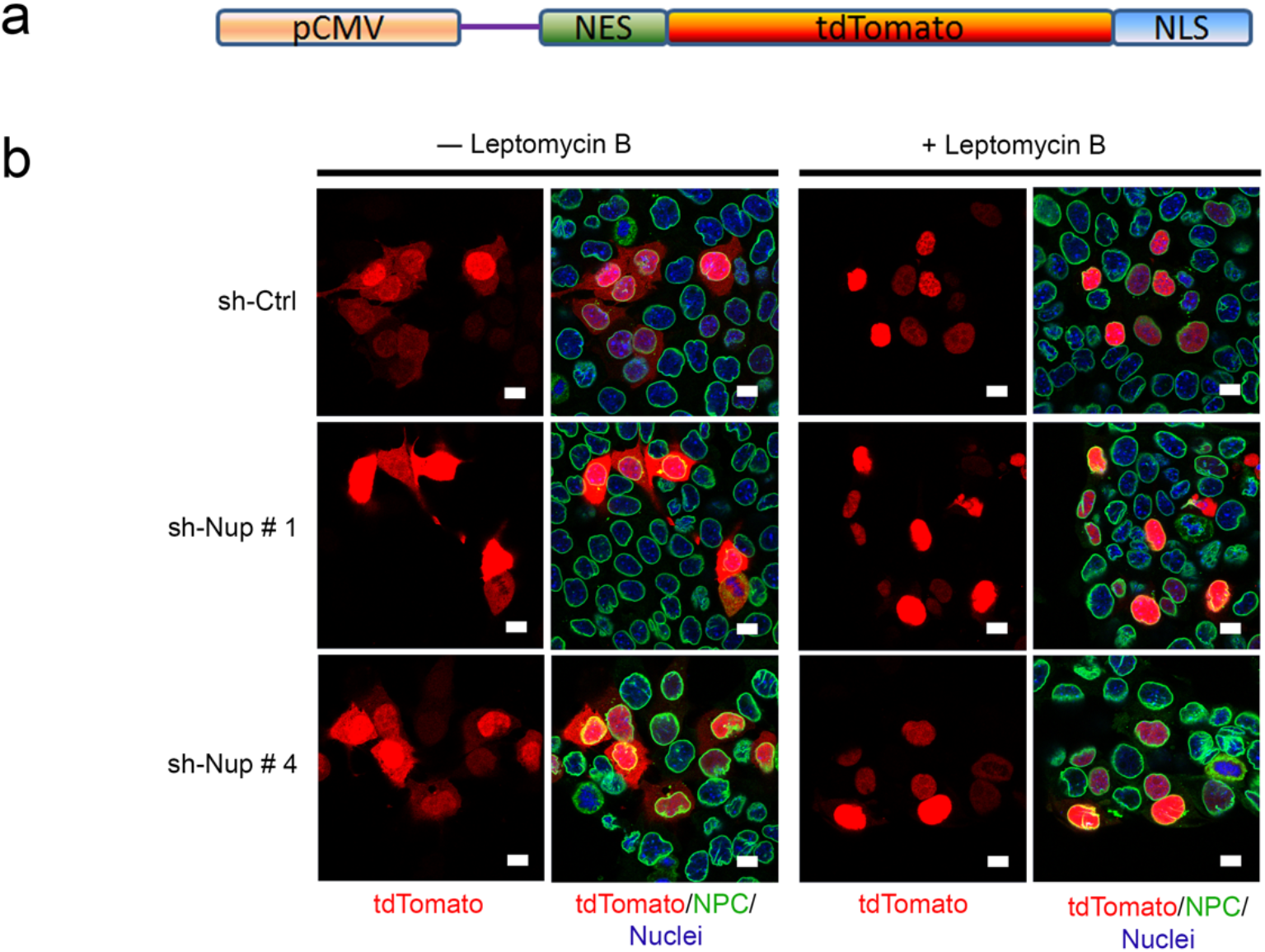
NUP210 loss does not affect general nucleocytoplasmic protein transport. (a) Schematic of the tdTomato-expressing nucleocytoplasmic transport reporter. Reporter expression is driven by a CMV promoter and produces tdTomato fused to both a nuclear export (NES) and import (NLS) signal. (b) Representative immunofluorescence images showing nuclear and cytoplasmic localization of the tdTomato signal in sh-Ctrl and *Nup210* KD 4T1 cells. The Nuclear Pore Complex (NPC) is shown in green. Nuclei are stained with DAPI. Leptomycin B, a nuclear export inhibitor drug, was used as positive control. Scale bar = 10 μm.

**Extended Data Figure 5:**
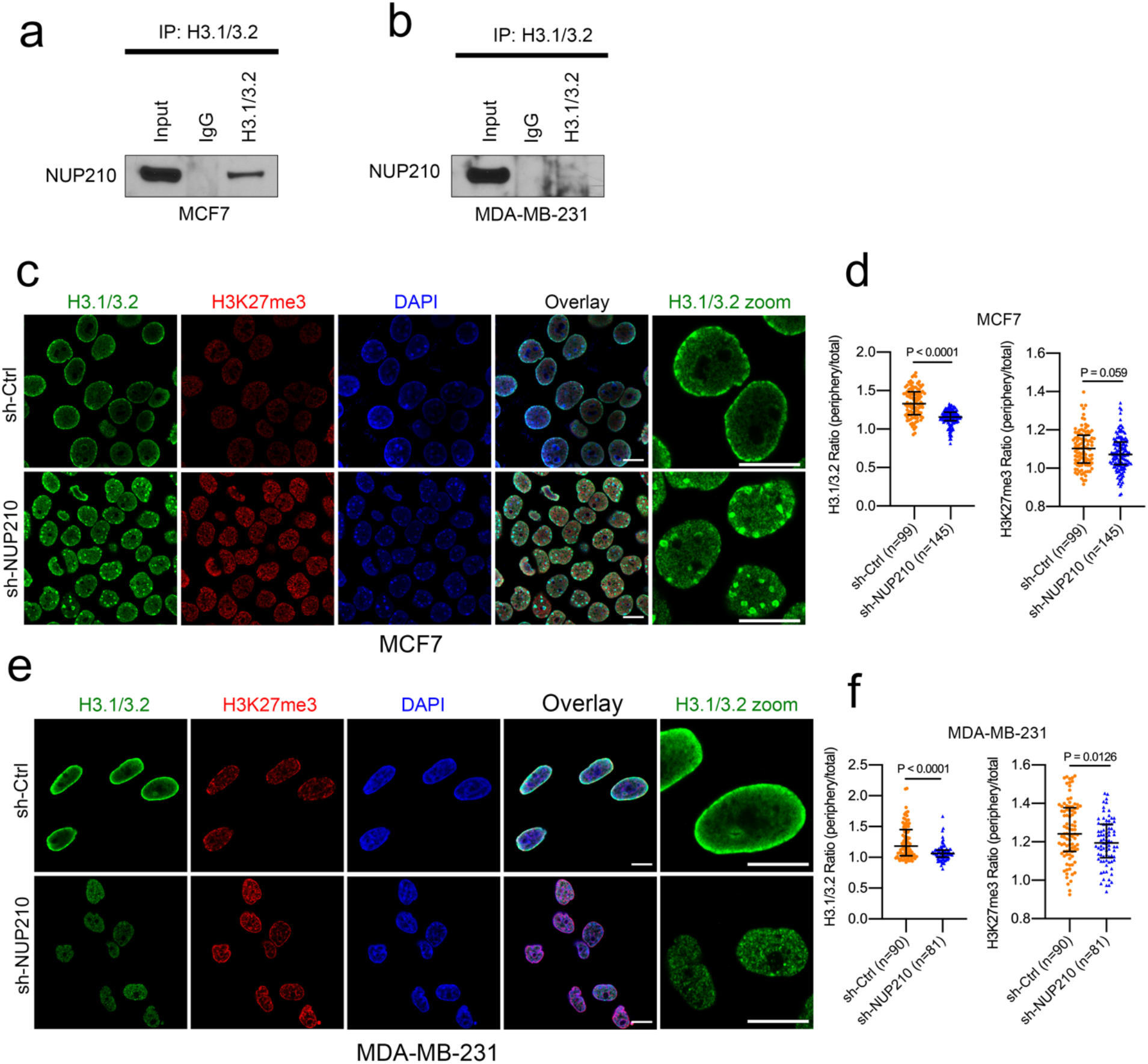
NUP210 interaction with H3.1/3.2 in human breast cancer cell lines MCF7 and MDA-MB-231. (a) Co-IP of NUP210 and H3.1/3.2 in MCF7 cell line. (b) Co-IP of NUP210 and H3.1/3.2 in MDA-MB-231 cell line. (c) Immunofluorescence staining of H3.1/3.2 and H3K27me3 in NUP210 knockdown MCF7 cells. Scale bar = 10 μm. (d) Quantification of H3.1/3.2 and H3K27me3 intensity mentioned in (c). Mann-Whitney U test, median with interquartile range. (e) Immunofluorescence staining of H3.1/3.2 and H3K27me3 in NUP210 knockdown MDA-MB-231 cells. Scale bar = 10 μm. (f) Quantification of H3.1/3.2 and H3K27me3 intensity mentioned in (e). Mann-Whitney U test, median with interquartile range.

**Extended Data Figure 6: Related to Figure 5.**
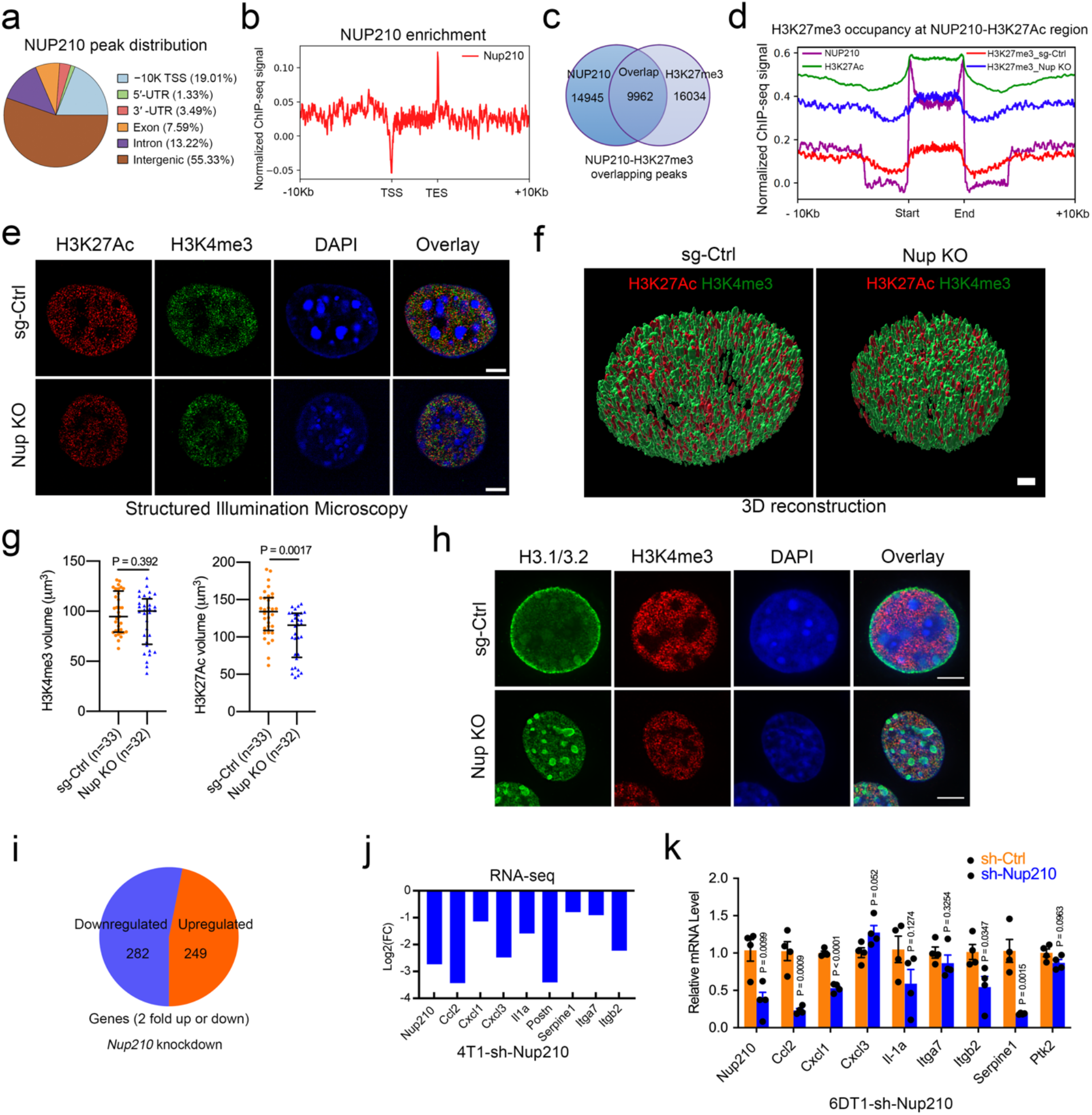
(a) Pie chart of NUP210 ChIP-seq peak distribution. (b) ChIP-seq profile of NUP210 enrichment. TSS = Transcriptional start site, TES = Transcriptional end site. (c) Overlap of NUP210 and H3K27me3 peaks. (d) Occupancy of H3K27me3 peaks within the NUP210-H3K27Ac enhancer overlap region in sg-Ctrl and *Nup210* KO cells. (e) Structured Illumination Microscopy images of H3K27Ac and H3K4me3 distribution in *Nup210* KO 4T1 cells. Scale bar = 5 μm. (f) 3D reconstruction of H3K27Ac and H3K4me3 volume distribution in *Nup210* KO 4T1 cells. Scale bar = 2 μm. (g) Quantification of H3K27Ac and H3K4me3 volume in *Nup210* KO 4T1 cells. Mann-Whitney U test, median with interquartile range. (h) Representative images of H3.1/3.2 and H3K4me3 distribution in *Nup210* KO 4T1 cells. Scale bar = 5 μm. (i) RNA-seq analysis of up- or downregulated genes upon *Nup210* KD. (j) Top downregulated genes from the RNA-seq of 4T1 *Nup210* KD (sh # 4) cells. FC=fold change. (k) qRT-PCR analysis on *Nup210* KD 6DT1 cells. Multiple two tailed t-test, mean ± s.e.m, n = 4.

**Extended Data Figure 7:**
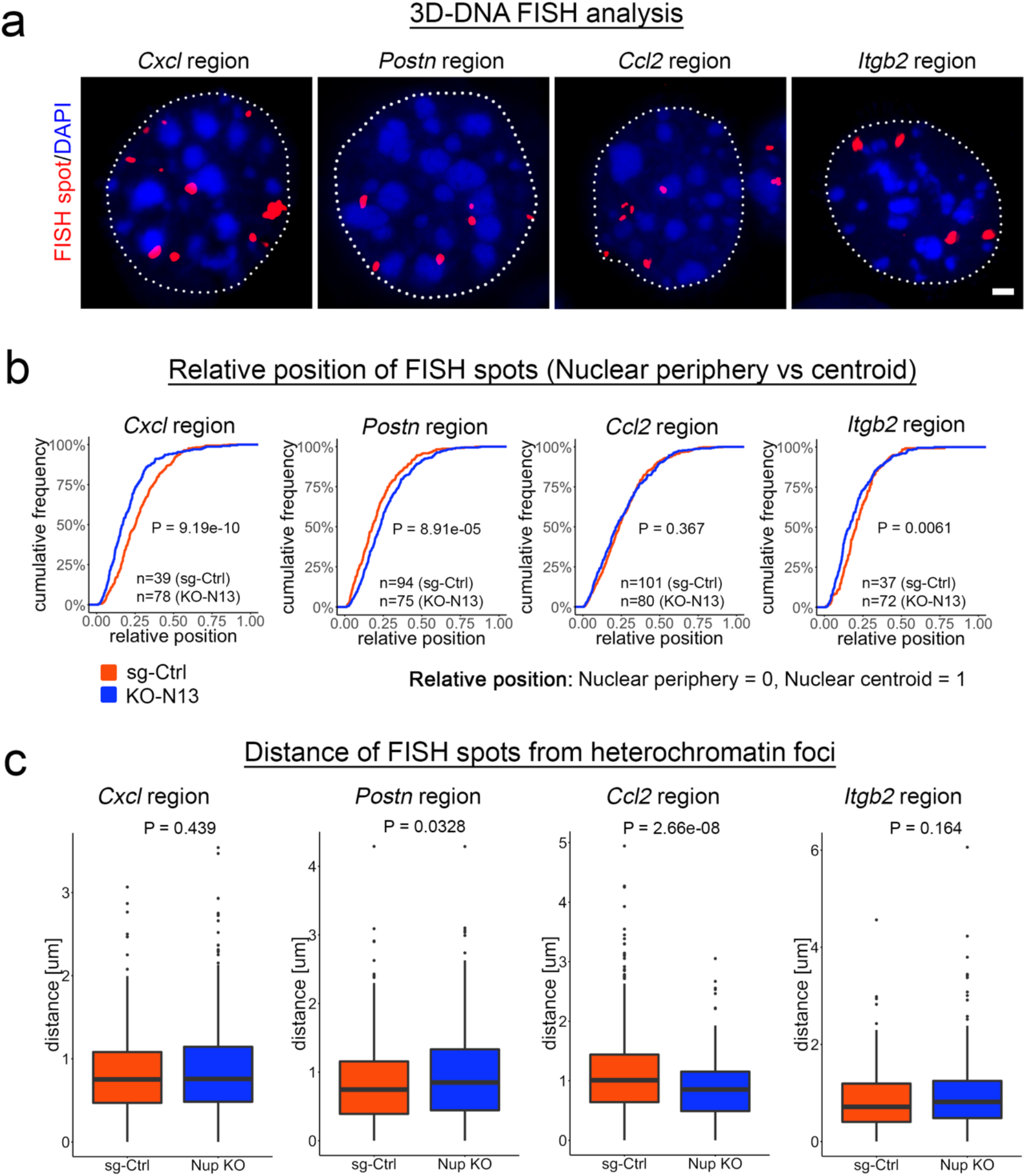
NUP210 loss is associated with differential repositioning mechanosensitive gene loci. (a) Representative images of 3D-DNA FISH of NUP210-regulated gene loci within the nucleus. Scale bar = 2 μm. (b) Cumulative distribution of FISH spots in nuclear periphery vs nuclear centroid in sg-Ctrl and *Nup210* KO 4T1 cells. (c) Minimum distance of FISH spots from DAPI-stained heterochromatin foci in sg-Ctrl and *Nup210* KO 4T1 cell nuclei stated in (b).

**Extended Data Figure 8: Related to Figure 6.**
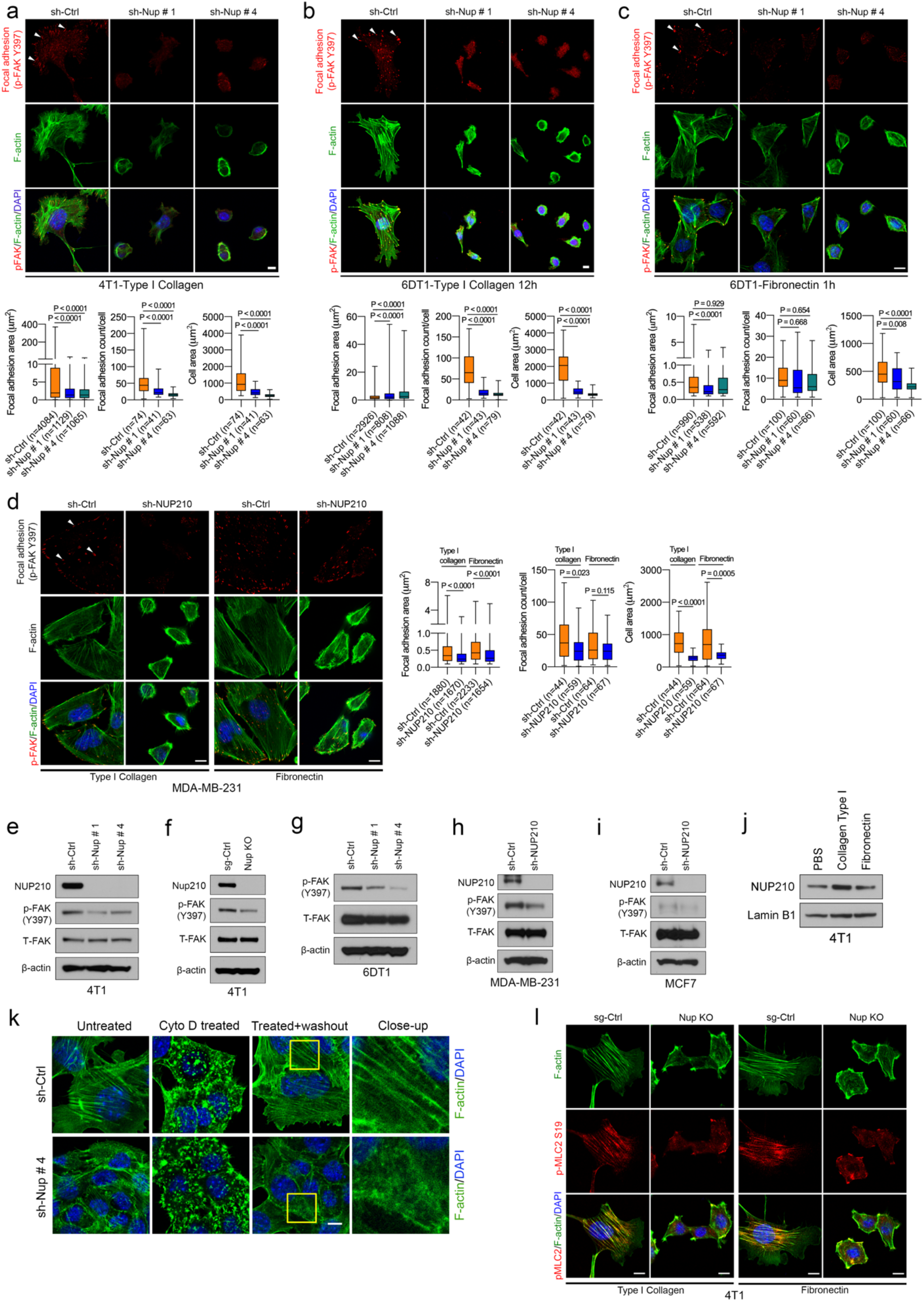
(a) (Top) Immunostaining of p-FAK Y397 and F-actin in *Nup210* KD 4T1 cells grown on Type I collagen. (Bottom) Quantification of focal adhesion (area, count) and cell spreading. ANOVA with Tukey’s multiple comparison test. Scale bar = 10 μm. (b) (Top) Immunostaining of p-FAK Y397 and F-actin in *Nup210* KD 6DT1 cells grown on Type I collagen. (Bottom) Quantification of focal adhesion (area, count) and cell spreading. ANOVA with Tukey’s multiple comparison test. Scale bar = 10 μm. (c) (Top) Immunostaining of p-FAK Y397 and F-actin in *Nup210* KD 6DT1 cells grown on fibronectin. (Bottom) Quantification of focal adhesion (area, count) and cell spreading. ANOVA with Tukey’s multiple comparison test. Scale bar = 10 μm. (d) (Left) Immunostaining of p-FAK Y397 and F-actin in *NUP210* KD MDA-MB-231 cells grown on Type I collagen or fibronectin. (Right) Quantification of focal adhesion (area, count) and cell spreading. ANOVA with Tukey’s multiple comparison test. Scale bar = 10 μm. (e) Western blot of total FAK (T-FAK) and p-FAK (Y397) levels in *Nup210* KD 4T1, (f) *Nup210* KO 4T1, (g) *Nup210* KD 6DT1, (h) *NUP210* KD MDA-MB-231 and (i) *NUP210* KD MCF7 cells. (j) Western blot of NUP210 and Lamin B1 on 4T1 cells grown on either Type I collagen or fibronectin. (k) Representative images of F-actin stress fibers (phalloidin) after treatment and washout of cytochalasin D in sh-Ctrl and *Nup210* KD 4T1 cells. Scale bar = 10 μm. (l) Representative images showing p-MLC2-S19 and F-actin staining on *Nup210* KO 4T1 cells grown on either Type I collagen or fibronectin. Scale bar = 10 μm. For the box plots, the box extends from 25th to 75th percentile, whiskers extend from smallest values to the largest values, horizontal line represents median.

**Extended Data Figure 9:**
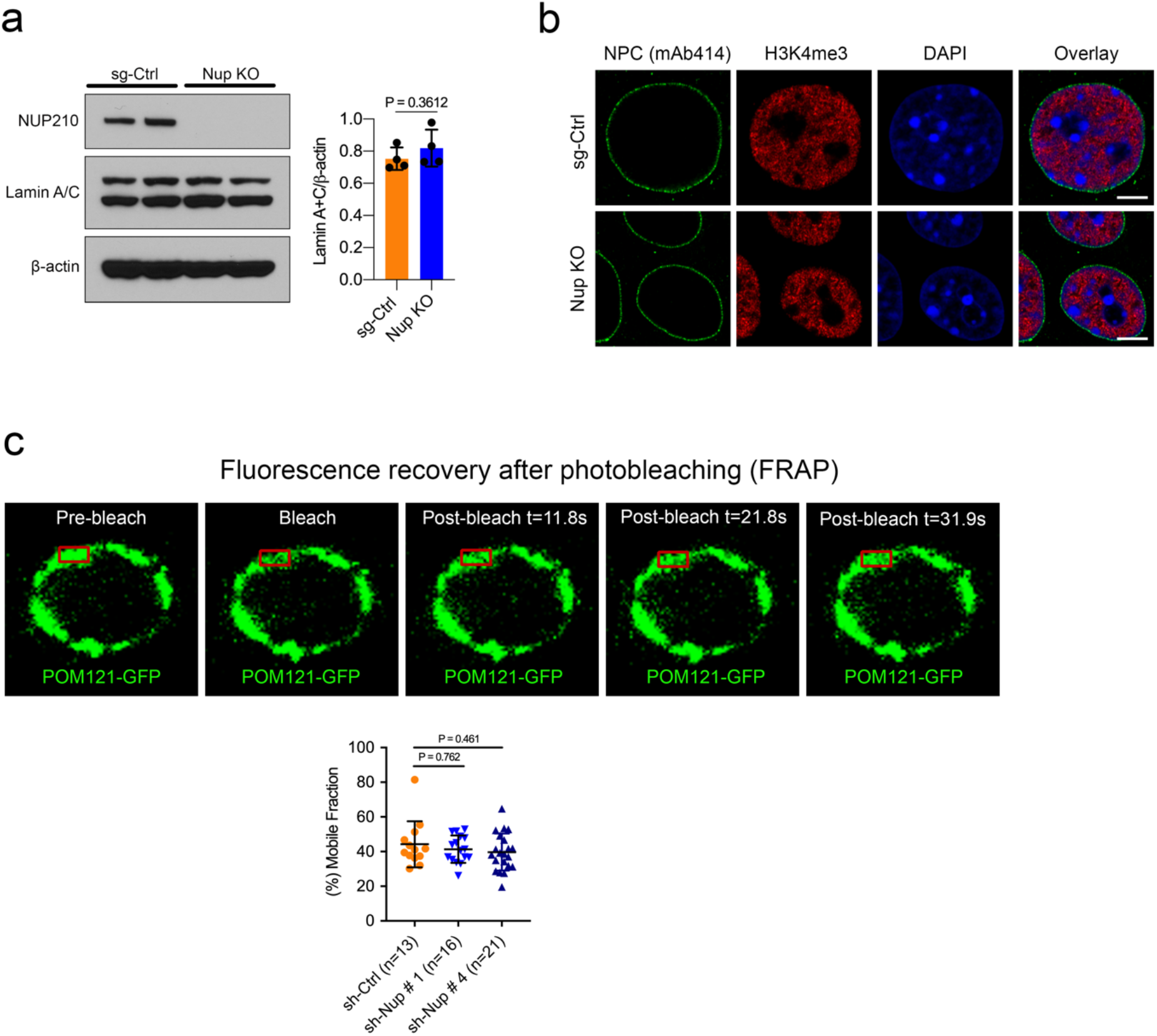
Effect of *Nup210* KO on Lamin A/C and on the dynamic distribution of nuclear pore structure. (a) Representative western blot showing the protein level of Lamin A/C (Left) and densitometry quantification of intensity normalized to *β*-actin (right). Two tailed t-test. Mean ± S.D. (b) Representative immunofluorescence images of the nuclear pore complex (stained using mAb414 antibody) and H3K4me3 in *Nup210* KO 4T1 cells. Scale bar = 5 μm. (c) (Top) Fluorescence Recovery after Photobleaching (FRAP) analysis showing the dynamic distribution of GFP-tagged nuclear pore protein, POM121 in *Nup210* KD 4T1 cells. (Bottom) Quantification of POM121-GFP mobile fraction. ANOVA with Tukey’s multiple comparison test, mean ± s.d.

**Extended Data Figure 10: Related to Figure 7.**
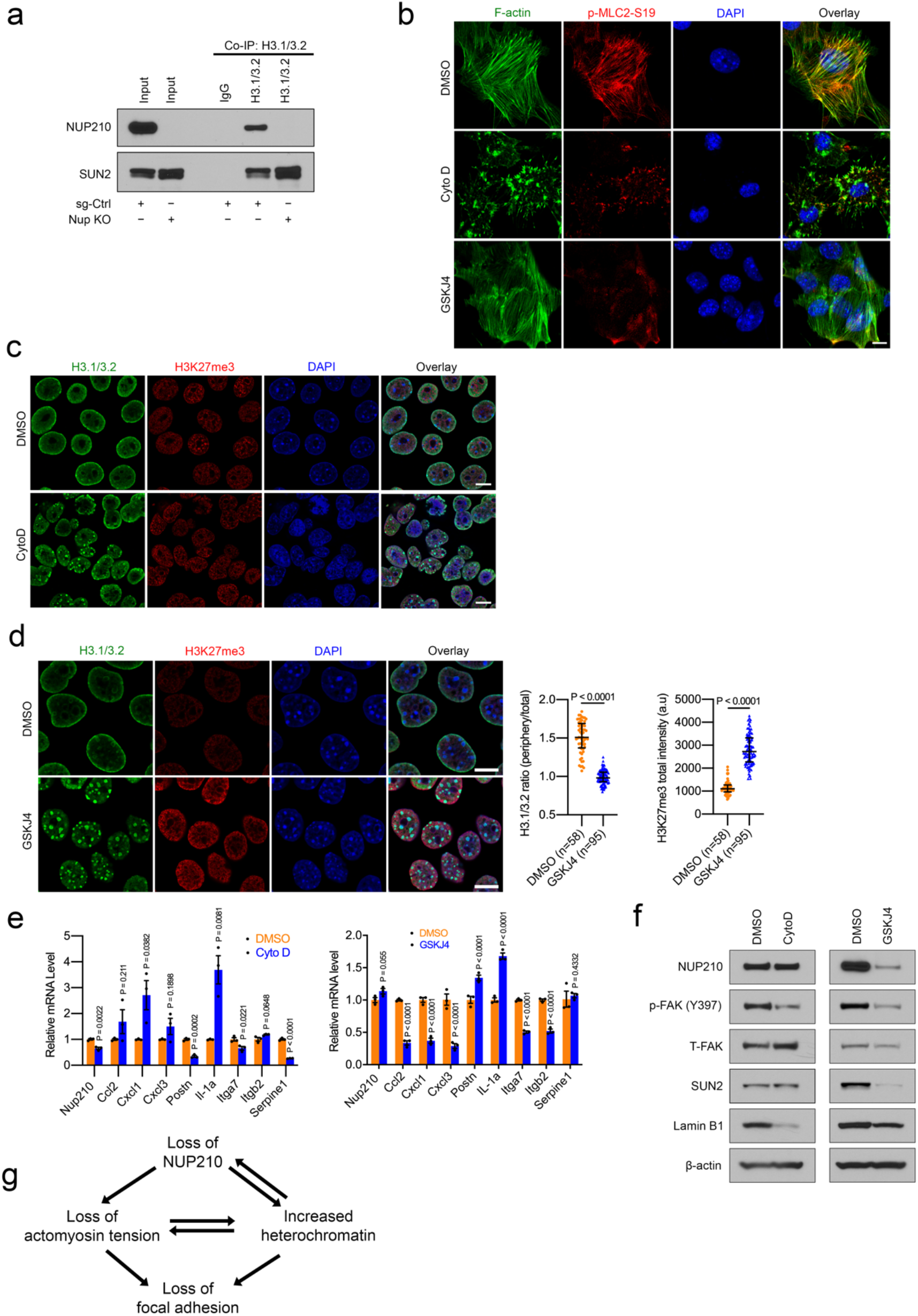
(a) Co-IP showing the interaction of H3.1/3.2 with NUP210 and SUN2 in *Nup210* KO 4T1 cells. (b) Representative immunofluorescence images of F-actin and p-MLC2-S19 staining in Cytochalasin D or GSKJ4-treated 4T1 cells. Scale bar = 10 μm. (c) Representative immunofluorescence images of H3.1/3.2 and H3K27me3 staining in Cytochalasin D-treated 4T1 cells. Scale bar = 10 μm. (d) (Left) Representative immunofluorescence images of H3.1/3.2 and H3K27me3 staining in GSKJ4-treated 4T1 cells. (Right) Quantification of intensity. Mann-Whitney U test, median with interquartile range. Scale bar = 10 μm. (e) (Left) qRT-PCR of NUP210-regulated genes in Cytochalasin D-treated and (right) GSKJ4-treated 4T1 cells. Multiple two tailed t-test, mean ± s.e.m. (f) Western blot of NUP210, p-FAK Y397, T-FAK, SUN2 and Lamin B1 in either Cytochalasin D- or GSKJ4-treated 4T1 cells. (g) Model depicting the feedback loop among the loss of NUP210, actomyosin tension, focal adhesion and and heterochromatin regulation.

### Supplementary Video files

**Supplementary Video 1:** 3D reconstruction of Z-stack images from H3.1/3.2 and H3K9me3 staining in 4T1 sg-Ctrl cells.

**Supplementary Video 2:** 3D reconstruction of Z-stack images from H3.1/3.2 and H3K9me3 staining in 4T1 *Nup210* KO cells.

**Supplementary Video 3:** 3D reconstruction of Z-stack images from H3K27Ac (enhancer) and H3K4me3 (promoter) staining in 4T1 sg-Ctrl cells.

**Supplementary Video 4:** 3D reconstruction of Z-stack images from H3K27Ac (enhancer) and H3K4me3 (promoter) staining in 4T1 *Nup210* KO cells.

**Supplementary Video 5:** Live cell imaging of F-tractin (red) and MLC2 (Cyan) reporter in 6DT1 sh-Ctrl cells after treatment with Cytochalasin D.

**Supplementary Video 6:** Live cell imaging of F-tractin (red) and MLC2 (Cyan) reporter in 6DT1 sh-*Nup210* cells after treatment with Cytochalasin D.

**Supplementary Video 7:** Live cell tracking of cell migration (measured by Hoechst staining of nuclei) in 4T1 sg-Ctrl cells.

**Supplementary Video 8:** Live cell tracking of cell migration (measured by Hoechst staining of nuclei) in 4T1 *Nup210* KO-N9 cells.

**Supplementary Video 9:** Live cell tracking of cell migration (measured by Hoechst staining of nuclei) in 4T1 *Nup210* KO-N13 cells.

**Supplementary Video 10:** 3D reconstruction of Z-stack images from Lamin B1 (red), Lamin A/C (green) and heterochromatin foci (DAPI, cyan) staining in 4T1 sg-Ctrl cells.

**Supplementary Video 11:** 3D reconstruction of Z-stack images from Lamin B1 (red), Lamin A/C (green) and heterochromatin foci (DAPI, cyan) staining in 4T1 *Nup210* KO cells.

## CONTACT FOR REAGENT AND RESOURCE SHARING

Further information and request for resources or reagents should be directed to and will be fulfilled by the corresponding authors, Ruhul Amin or Kent W. Hunter.

## METHODS

### Mouse strains

Usage of animals described in this study was performed under the animal study protocol LCBG-004 approved by the National Cancer Institute (NCI) at Bethesda Animal Use and Care Committee. Animal euthanasia was performed by anesthesia using Avertin injection followed by cervical dislocation. Female BALB/c (000651) and FVB/NJ (001800) mice were purchased from The Jackson Laboratory.

### Cell lines

Mouse mammary tumor cell lines, 4T07, 4T1, 6DT1, and MVT1, were provided by Dr. Lalage Wakefield (NCI, NIH)^26^. These cell lines were grown in DMEM (Gibco) supplemented with 9% fetal bovine serum (FBS) (Gemini), 1% L-glutamine (Gibco), and 1% penicillin-streptomycin (Gemini). Human breast cancer cell lines, MDA-MB-231 and MCF7, were provided by Dr. Jeffrey E. Green (NCI, NIH) and grown in DMEM with 9% FBS, 1% L-glutamine, and 1% Penicillin-Streptomycin. Human 293FT cells were purchased from Thermo Fisher Scientific and grown on the medium mentioned above.

### Benzonase Accessible Chromatin (BACh) sequencing analysis

Cells were expanded in DMEM at 37°C to obtain 8–10 x10^7^ cells/condition/each experiment. BACh analysis was performed as previously described with minor modifications ^75^. Briefly, cells were collected by centrifugation, washed twice in ice cold cellular wash buffer (20 mM Tris-HCl pH 7.5, 137 mM NaCl, 1 mM EDTA, 10 mM sodium butyrate, 10 mM sodium orthovanadate, 2 mM sodium fluoride, protease inhibitor cocktail (Roche) and resuspended in (40 million cells/ml) hypertonic lysis buffer (20 mM Tris-HCl pH 7.5, 2 mM EDTA, 1 mM EGTA, 0.5% glycerol, 20 mM sodium butyrate, 2 mM sodium orthovanadate, 4 mM sodium fluoride and protease inhibitor cocktail). Cells were distributed in 500 µl aliquots in 1.5 ml tubes and followed by the addition of 500 µl of nuclease digestion buffer (40 mM Tris-HCl pH 8.0, 6 mM MgCl_2_, 0.3% NP-40, and 1% Glycerol) containing a 3-fold dilutions (from 0.125 units/ml to 6 units/ml) of Benzonase nuclease (Millipore). This was mixed gently and incubated for three minutes at 37°C. Reactions were terminated by the addition of EDTA (10 mM final concentration) and SDS (0.75% final concentration). Proteinase K was added to a final concentration of 0.5 mg/ml and incubated overnight at 45°C. DNA fragments of 100-500 bp from a chromatin digestion were purified over sucrose gradients^76^ and precipitated in 0.1 volume sodium acetate and 0.7 volume isopropanol.

### Sequencing and data analysis of Benzonase-treated samples

DNA was sequenced using either Illumina HiSeq2000 (TrueSeq V3 chemistry) or NextSeq500 (TrueSeq High output V2 chemistry) sequencers at the Advanced Technology Research Facility (ATRF), National Cancer Institute (NCI-Frederick, MD, USA). The sequence reads were generated as either 50-mer or 75-mer (trimmed to 50-mer by trimmomatic software before alignment), and tags are then aligned to the UCSC mm9 reference genome assembly using Eland or Bowtie2. All the samples were in good quality with over 94% of the bases having Q30 or above with 25∼50 million raw reads per sample. Regions of enriched tags termed hotspots have been called using DNase2Hotspots algorithm^77^ with FDR of 0% with minor updates. Tag density values were normalized to 10 million reads. For comparison of BACh profiles across multiple cell lines, Percent Reference peak Coverage (PRC) measure was proposed and employed to ensure compatible levels of digestion by Benzonase in multiple cell lines^78^. To calibrate PRC, commonly represented hotspot sites were identified as reference peaks based on mouse (mm9) DNaseq data from ENCODE. ENCODE narrowPeak definition files (DNaseI Hypersensitivity by Digital DNaseI from ENCODE/University of Washington) for a total of 133 samples were downloaded from the UCSC golden path web site: http://hgdownload.cse.ucsc.edu/goldenPath/mm9/encodeDCC/wgEncodeUwDnase/. A total of 8,587 peaks were found to be present in all 133 samples. By assuming that the most commonly accessible sites may also be present in our optimally digested samples, PRC was obtained as a fraction of reference peaks presented in each sample to evaluate the level of digestion by Benzonase. Our pooled samples with PRC of 90% or greater indicated that all the 8,587 sites were present in the sample. Two biological replicates with acceptable PRC were selected and pooled for each cell line, and each pooled data set showed over 90% PRC. When samples showed bimodality in their tag density distribution due to elevated noise, we dropped hotspots that belonged to a group with low tag density and low PRC by applying threshold to maximum hotspot tag density values.

### Computing environment

All computations at NCI were performed on the NIH helix/biowulf system, documentation of which is available at https://helix.nih.gov. We used the R computing environment, Perl scripts, Bedtools, and UCSC liftOver for most of the analyses.

### Identification of polymorphic BACh sites

The workflow consisted of the following: 1) the BACh data were filtered for the regions overlapping with polymorphic sites. Since the BACh data were generated in Genome Build mm9, we used UCSC mm9 snp128 data to restrict the BACh sites. 2) Variant Called Format (VCF) files were filtered to retain the SNPs that overlap with the BACh present in the 4T1 and 4T07 cell lines. 3) SNPs were removed in the BACh that are present in the mouse FVB/NJ strain.

### Long Read Chromatin Interaction Analysis Through Paired-end Tag (ChIA-PET) sequencing and data processing

Long read ChIA-PET was performed using Tn5 transposase to tag DNA for long tag sequencing by Illumina NextSeq. ChIA-PET data was processed by a customized ChIA-PET data processing pipeline^22^. Detailed protocol can be found in Li et al.^79^.

### Cloning

A mouse *Nup210* full length cDNA (NM_018815) encoding vector (pCMV6-Nup210-Myc) was purchased from Origene Technologies. This vector was digested with SalI-HF (New England Biolabs) and EcoRV-HF (New England Biolabs) restriction enzymes to obtain the NUP210 protein-encoding region. This NUP210 insert was then cloned into the Gateway entry vector pENTR1A using the Quick Ligation Kit (New England Biolabs). pENTR1A no ccdB plasmid (Addgene plasmid # 17398) was received by Dr. Marian Darkin (NCI) as a gift from Dr. Eric Campeau. The NUP210-encoding region from the pCMV6-NUP210-Myc vector was cloned into pENTR1A vector through digestion with SalI and EcoRV. pENTR1A-NUP210 entry vector was used to transfer the NUP210-encoding region into the lentiviral destination vector pDEST-658 (received as a gift from Dr. Dominic Esposito, NIH) along with a mouse *Pol2* promoter entry vector using the Gateway LR clonase reaction (Thermo Fisher Scientific). The integrity of the final NUP210-encoding vector was verified through DNA sequencing.

3x Flag-Histone H3.1 plasmid was received as a gift from Dr. Jing Huang (NCI, NIH). Coding region of H3.1 along with Flag-Tag was amplified by PCR using KOD Hot Start DNA Polymerase (Millipore) and cloned into gateway entry vector pENTR/D-TOPO vector (Thermo Fisher Scientific). Entry vectors containing Flag-H3.1 was then cloned into gateway lentiviral destination vector pDest-659 (gift from Dr. Dominic Esposito Lab) along with mPol2 promoter and C-terminal V5 tag entry vectors using the gateway LR clonase reaction (Thermo Fisher Scientific).

The pGL4.23 [*luc2*/minP] promoter luciferase vector (Promega, Catalog # E8411) was received as a gift from Dr. Jing Huang (NCI, NIH). A 550 bp region from the mouse (FVB/NJ and BALB/cJ) *Nup210* promoter covering the polymorphic sites was amplified by PCR using KOD Hot Start DNA Polymerase (Millipore) and digested with KpnI (New England Biolabs) and XhoI (New England Biolabs) restriction enzymes. Gel-purified PCR product was then cloned into the pGL4.23 vector using T4 DNA ligase (New England Biolabs). Finally, the DNA sequence of the clones were confirmed through DNA sequencing.

pMXs-puro-EGFP-FAK plasmid was obtained as a gift from Dr. Noboru Mizushima (University of Tokyo) (addgene # 38194). cDNA encoding mouse FAK (Ptk2) without EGFP was PCR amplified from this plasmid and subcloned into pENTR/D-TOPO entry vector (Thermo Fisher Scientific). Entry vector is then further cloned into lentiviral destination vector pDest-659 along with mPol2 promoter and C-terminal myc-tag entry vectors using the method described above. cDNA encoding mouse *Ccl2* was PCR amplified from 4T1 cells and subcloned into pENTR/D-TOPO entry vector. Ccl2 entry vector is then further cloned into lentiviral destination vector pDest-659 along with mPol2 promoter and C-terminal myc-tag entry vectors using above method. Sequence of the final clones were confirmed through sanger sequencing.

### Lentivirus production and generation of stable cell lines

All of the TRC lentiviral shRNA vectors were purchased from Dharmacon. shRNAs targeting the mouse *Nup210* gene, sh-Nup # 1 (TRCN0000101935: TAACTATCACAGTAAGAAGGC) and sh-Nup # 4 (TRCN0000101938: TTCAGTTGCTTATCTGTCAGC) were used for *Nup210* knockdown in all of the mouse cell lines. For mouse *Ctcf* knockdown, sh-Ctcf # 1 (TRCN0000039022: TAAGGTGTGACATATCATCGG) and sh-Ctcf # 2 (TRCN0000039023: ATCTTCGACCTGAATGATGGC) were used. For mouse *Ccl2* knockdown, sh-Ccl2 # 3 (TRCN0000034471: TTACGGGTCAACTTCACATTC), sh-Ccl2 # 4 (TRCN0000034472: TTGCTGGTGAATGAGTAGCAG), and sh-Ccl2 # 5 (TRCN0000034473: AATGTATGTCTGGACCCATTC were used. For *NUP210* knockdown in human cell lines, a shRNA targeting the human *NUP210* gene (TRCN0000156619: AAATGAGCTAATGGGCAGAGC) was used.

For lentivirus production, shRNA-containing plasmids and packaging plasmids, psPAX2 (Addgene plasmid # 12260) and envelope plasmid pMD2.G (Addgene plasmid # 12259) (both were a gift from the Trono lab), were transfected into the human 293FT (Thermo Fisher Scientific) cell line using X-tremeGENE 9 DNA transfection reagent (Roche). 48 h after transfection, culture supernatant containing lentivirus was harvested, filtered through a 0.45 μm filiter (Millipore), and then used for transduction of mouse and human breast cancer cell lines. shRNAs stably integrated in mouse and human cells were selected with 10 and 2 μg/ml puromycin (Sigma), respectively. For the selection of NUP210-overexpressing cells, 10 μg/ml of blasticidin (Gibco) was used.

### CRISPR/Cas9-mediated knockout of the mouse Nup210 gene

For CRISPR/Cas9-mediated knockout of mouse *Nup210* in the 4T1 cell line, sgRNA targeting *Nup210* exon 5 (sgRNA sequence: GCGACACCATCCTAGTGTCT) was designed using the GPP Web Portal available at the Broad Institute (https://portals.broadinstitute.org/gpp/public/analysis-tools/sgrna-design). sgRNA was then cloned into the lentiGuide-puro (Addgene plasmid # 52963, a gift from the Feng Zhang lab) vector. Non-targeting control sgRNA cloned into the lentiGuide-puro vector (sgRNA sequence: CCATATCGGGGCGAGACATG) was kind of a gift from Dr. Ji Luo (NCI, NIH). Lentiviral particles were prepared as described above for shRNA lentiviruses. 4T1 cells stably expressing the sgRNAs were generated through lentiviral transduction and selection with 10 μg/ml puromycin. For transient expression of Cas9 in the sgRNA-stable 4T1 cells, an adenoviral Cas9-encoding viral particles containing the GFP reporter (Vector Biolab) was used at 25 M.O.I. 96 h after transfection of the Cas9 vector, GFP-positive 4T1 cells were FACS sorted and single cells were isolated for clonal expansion. *Nup210* mutation was confirmed through DNA sequencing and knockout was verified through western blot. Before performing functional assays with *Nup210* knockout cells, both sgCtrl and knockout cells were passaged at least 4 times for 2 weeks to eliminate residual Cas9-GFP signal within 4T1 cells, which was verified using a fluorescence microscope.

### CRISPR/Cas9 D10A-mediated deletion of CTCF-binding site on Nup210 promoter

For the deletion of polymorphic CTCF binding region on mouse *Nup210* promoter, Cas9 D10A double nicking strategy was used. Sense and antisense sgRNAs spanning the CTCF binding site were designed using the above link of GPP web portal of Broad Institute. All-in-One plasmid encoding dual U6 promoter-driven sgRNAs and EGFP-coupled Cas9-D10A plasmid (AIO-GFP) was a gift from Dr. Steve Jackson (The Wellcome Trust Sanger Institute, UK) (Addgene # 74119). sgRNAs were cloned into AIO-GFP plasmid using the method described in the paper^80^. Briefly, AIO-GFP plasmid was digested with *BbsI* restriction enzyme, dephosphorylated using calf intestinal phosphatase and gel purified. sgRNA oligonucleotide pairs were purchased from Integrated DNA Technologies, annealed and phosphorylated using T4 polynucleotide kinase (NEB). First sgRNA was cloned into *BbsI* site and second sgRNA was cloned into *BsaI* site. DNA sequence of the final clone was verified via sanger sequencing using the primer (5*′*-CTTGATGTACTGCCAAGTGGGC-3*′*). sgRNA plasmid was then transfected into 4T1 cells using Nanojuice transfection reagent (Millipore). 48 h later, GFP positive cells was sorted using flow cytometry and single cells were plated on 96 well plate. Cas9 D10A-edited clones were identified using the PCR and DNA sanger sequencing.

### Spontaneous metastasis assay in mice

6-8 weeks old female BALB/c and FVB/NJ mice were purchased from The Jackson Laboratory. For orthotopic transplantation of gene knockdown or overexpressing cells, 100,000 cells were injected into the fourth mammary fad pad of mice. 28-30 days after injection, mice were euthanized and primary tumors were resected, weighed, and surface lung metastases were counted.

### Protein nucleocytoplasmic transport assay

4T1 *Nup210* knockdown cells were grown on glass coverslips in 2-well Lab-Tek chambered glass coverslip (Thermo Fisher Scientific) at a seeding density of 20,000 cells/well. 24 h after seeding, cells were transfected with 1 μg of nucleocytoplasmic transport reporter (NLS-tdTomato-NES) plasmid (received as a gift from Dr. Martin W. Hetzer, Salk Institute) using Novagen Nanojuice Transfection Reagent (Millipore-Sigma). 4 h after transfection, medium was replaced with fresh medium and cells were grown for 24 h. Cells were then treated with 20 nM leptomycin B (Cell Signaling Technology) for 6 h and then fixed with methanol for immunostaining with an antibody against nuclear pore complex proteins (clone mAb414, Abcam) following the regular immunofluorescence protocol described below. Nuclear and cytoplasmic expression of tdTomato was observed using confocal microscopy.

### Protein interaction analysis through Liquid Chromatography-Mass Spectrometry (LC-MS)

NUP210-Myc overexpressing 4T1 cells were seeded onto 15 cm tissue culture dishes at a seeding density of 2.5×10^6^ cells/dish. After 48 h of incubation, cells were harvested and nuclear protein complex lysates were prepared using the Nuclear Complex Co-IP Kit (Active Motif). Co-immunoprecipitation with two biological replicates was performed according to the manufacturer’s instructions. Briefly, 500 μg of nuclear protein lysates were incubated with 2 μg of either Myc-Tag antibody (Cell Signaling Technology) or an endogenous NUP210-specific antibody (Bethyl Laboratories). After overnight incubation on a rotator at 4°C, 25 μg of Dynabeads Protein G (Invitrogen) were added to the protein lysate-antibody complexes. After 30 min of incubation with beads, antibody-bead-protein complexes were isolated using a magnetic stand. Beads were then washed three times and then dissolved in 25 mM ammonium bicarbonate pH 8.0 (Sigma). Samples were then subjected to LC-MS analysis.

### Western blot

Whole cell protein lysate was prepared using lysis buffer (20 mM Tris-HCl pH 8.0, 400 mM NaCl, 5 mM EDTA, 1 mM EGTA, 10 mM NaF, 1 mM sodium pyrophosphate, 1% Triton X-100, 10% glycerol, protease and phosphatase inhibitor cocktail). Nuclear and cytoplasmic protein lysates were prepared using the Nuclear Extract Kit (Active Motif) or Nuclear Complex Co-IP Kit (Active Motif). Protein concentration was measured using the Pierce BCA Protein Assay Kit (Thermo Fisher Scientific). 25 μg of protein lysates were mixed with 4x NuPAGE LDS sample buffer (Invitrogen) and 10x NuPAGE Sample Reducing Agent (Invitrogen). Samples were then boiled at 95°C for 5 min and resolved on NuPAGE 3-8% Tris-acetate, NuPAGE 4-12% Bis-Tris, or Novex 4-20% Tris-Glycine protein gels (Thermo Fisher Scientific) with appropriate running buffer. Protein was transferred onto a PVDF membrane (Millipore) and the membrane was blocked with blocking buffer (TBST + 5% Non-fat dry milk) for 1 h. Membranes were then incubated with appropriate primary antibodies overnight. After washing with TBST, membranes were incubated with secondary antibodies for 1 h. Finally, the signal was developed on X-ray film using the Amersham ECL Western Blotting Detection Reagent (GE Healthcare). Densitometry quantification of western blot was performed using FIJI software^81^.

Primary antibodies and their dilutions were as follows: NUP210 (1:500; Bethyl Laboratories), *β*-actin (1:10,000; Abcam), Lamin B1 (1:5,000; Abcam), Lamin A/C (1:1,000; Abcam), Myc-Tag (1:1,000; Cell Signaling Technology), rabbit V5-Tag (1:1,000; Cell Signaling Technology), mouse V5-Tag (1:1,000; Cell Signaling Technology), RRP1B (1:1,000; Millipore-Sigma), SUN1 (1:500; Abcam), SUN2 (1:1,000; Abcam), FAK (1:5,000; Abcam), p-FAK(Y397) (1: 5,000; Abcam), p-FAK (Y397) (1:5,000; Thermo Fisher Scientific), CCL2 (1:1,000; Proteintech), H3.1/3.2 (1:1,000; Active Motif), H3K27me3 (1:5,000; Cell Signaling Technology), H3K9me3 (1:5,000; Abcam), SUV39H1 (1:1,000, Cell Signaling Technology), EZH2 (1:1,000; Cell Signaling Technology), and SUZ12 (1:1,000; Cell Signaling Technology). Anti-mouse secondary antibody (GE Healthcare) was used at 1:10,000 dilution and anti-rabbit secondary antibody (Cell Signaling Technology) was used at 1:3,000 dilution.

### Co-immunoprecipitation

Co-immunoprecipitiation was performed using the Nuclear Complex Co-IP Kit (Active Motif). 4T1 cells were seeded onto 15 cm tissue culture dishes at a seeding density of 5×10^6^ cells/dish. After 48 h of incubation, cells were harvested and nuclear lysates were prepared. 200-500 μg of nuclear lysates were incubated with 2 μg of specific antibodies and 50 μg of Dynabeads Protein G (Invitrogen). After overnight incubation on a rotator at 4°C, immune complexes were isolated using a magnetic stand. Beads were then washed three times, resuspended in 2x NuPAGE LDS sample buffer (Invitrogen), and incubated at 95°C heat block for 5 min. Samples were loaded onto NuPAGE protein gels and the standard western blot protocol was followed as described above.

For co-immunoprecipitation in 293FT cells, 2.5×10^6^ cells were seeded into 15 cm tissue culture dishes. Flag-H3.1 (5 μg) and NUP210-Myc (5 μg) plasmids were co-transfected using Xtremegene 9 transfection reagent. 48h after the transfection, cells were harvested and nuclear protein lysates were prepared using the Nuclear Complex Co-IP kit (Activemotif). Co-immunoprecipitation was followed as described above.

### Immunofluorescence and confocal microscopy

Immunofluorescence analysis was performed as described previously^28^. Briefly, cells were grown on 4-well or 8-well polymer coverslips (Ibidi) at a seeding density of 40,000 or 20,000 cells/well, respectively. After 24 h of incubation, cells were fixed with −20°C methanol for 2 min and permeabilizaed with PBS containing 1% Triton X-100 for 1 min. Fixed cells were then blocked with immunofluorescence buffer (1x PBS, 10 mg/ml BSA, 0.02% SDS, and 0.1% Triton X-100) for 30 min. Cells were then incubated with primary antibodies diluted in immunofluorescence buffer overnight at 4°C. After washing the cells with immunofluorescence buffer three times for 10 min per wash, cells were incubated with Alexa Fluor-conjugated secondary antibodies for 1 h at room temperature. Cells were washed three times with immunofluorescence buffer and then incubated with 1 μg/ml DAPI (4′,6-diamidino-2-phenylindole) for 10 min to stain the nucleus. After washing the cells with PBS three times, slides were kept at 4°C until subjected to confocal microscopy. In cases of drug treatment followed by super-resolution microscopy, cells were treated with 1 μM JQ1, 5 μM GSKJ4 and 0.5 μM Cytochalasin D for 24h. Images were acquired using either a Zeiss LSM 780 confocal microscope (63x plan-apochromat N.A. 1.4 oil immersion objective lens, 0.09 μm X-Y pixel size and 1.0 μm optical slice thickness), a Zeiss LSM 880 Airyscan super-resolution microscope (Airyscan detector, 63x plan-apochromat N.A. 1.4 oil immersion objective lens and 0.05 μm X-Y pixel size), a Zeiss LSM 780 Elyra with Structured Illumination Microscopy (SIM) module or a Nikon SoRa spinning disk super-resolution microscope (Yokogawa SoRa spinning disk unit, 60x plan-apochromat N.A. 1.49 oil immersion objective lens, Photometrics BSI sCMOS camera, and 0.027 μm X-Y pixel size). Airyscan images were processed using the Airyscan processing algorithm in the Zeiss ZEN Black (v.2.3) software, whereas the Nikon SoRa images were deconvolved using a constrained iterative restoration algorithm in the Nikon NIS Elements (v5.11) software. Tetraspeck 0.2 μm beads (Invitrogen) were imaged with the same microscope parameters and used for channel alignment.

Primary antibodies used for immunofluorescence were as follows: p-FAK (Y397) (1:100; Abcam), H3.1/3.2 (1:1,000; Active Motif), mouse Myc-Tag (1:500; Cell Signaling Technology), rabbit Myc-Tag (1:100; Cell Signaling Technology), SUN1 (1:100; Abcam), SUN2 (1:100; Millipore), p-MLC2-S19 (1:100; Cell Signaling Technology), p-FAK Y397 (1:500 for tissue IF; Thermo Fisher Scientific), H3K4me3 (1:1,000; Millipore), rabbit H3K27me3 (1:1,000; Cell Signaling Technology), mouse H3K27me3 (1:250; Abcam), H3K27Ac (1:500; Cell Signaling Technology), H3K9me3 (1:1,000; Abcam), Nucleolin (1:500; Abcam), Lamin B1 (1:500; Abcam), Lamin A/C (1:500; Abcam) and Nuclear Pore Complex (NPC) antibody (clone mAb414, 1:1,000; Abcam). Secondary antibodies used were mouse Alexa Fluor 488 (1:200; Invitrogen), Phalloidin Alexa Fluor 488 (1:250; Invitrogen), rabbit Alexa Fluor 488 (1:200; Invitrogen), rabbit Alexa Fluor 568 (1:200; Invitrogen), rabbit Alexa Fluor 594 (1:200; Invitrogen), and mouse Alexa Fluor 594 (1:200).

### Immunofluorescence on tumor tissue

Frozen tumor tissue sections were fixed in 4% paraformaldehyde in PBS for 10 minutes at room temperature. Slides were then washed with PBS 3 times and permeabilized with 0.5% Triton X-100 in PBS for 3 minutes. After washing with 1X PBS 3 times, slides were then incubated with blocking solution (10% BSA in PBS) for 1 hour. Primary antibodies were diluted in blocking solution, added to the tissue sections and incubated overnight at 4°C. Slides were washed 3 times with PBS + 0.1% Tween 20 (PBST) for 5 minutes and incubated with secondary antibodies in blocking buffer for 1 hour at room temperature. Sections were then washed with 1X PBST 3 times for 5 minutes and incubated with DAPI solution (1 μg/ml) for 10 minutes at room temperature. Slides were washed with 1X PBS 3 times for 5 minutes and then mounted with Prolong Glass Antifade mountant (Invitrogen). Confocal Z-stack images were acquired using Nikon SoRA spinning disk microscope. Dilution of the primary antibodies are as follows: rabbit p-FAK Y397 (1:500; Thermo Fisher Scientific), mouse alexa Fluor 488-conjugated pan-Cytokeratin (1:100; Cell Signaling Technology). Dilution of the secondary antibodies are as follows: Anti-rabbit Alexa Fluor 568 antibody (1:200; Invitrogen).

### Fluorescence Recovery After Photobleaching (FRAP) Assay

4T1 cells were grown on 35mm glass-bottom dishes (MatTek). After overnight incubation, cells were transfected with 0.5 μg POM121-GFP plasmid (Genecopoeia) using Nanojuice transfection reagent. 24h after the transfection, cells were subjected to FRAP analysis in Zeiss 880 confocal microscope. Percent mobile fraction of POM121-GFP in each condition was calculated using the FRAP module fit formula option of the ZEN software (Zeiss).

### Cell cycle analysis

Cell cycle analysis was performed using the Click-iT EdU Alexa Fluor 488 Flow Cytometry Assay Kit (Thermo Fisher Scientific) and FxCycle Violet Stain (Thermo Fisher Scientific) according to manufacturer’s instructions. Briefly, 4T1 cells were seeded onto 15 cm tissue culture dishes at a seeding density of 3×10^6^ cells/dish. 24 h later, cells were pulsed with Click-iT EdU (5-ethynyl-2’-deoxyuridine) for 1 h. After harvesting the cells through trypsinization, cells were fixed with Click-iT fixative and permeabilized with saponin-based permeabilization agent. The Click-iT reaction was then performed for 30 min at room temperature. Cells were then washed with wash buffer and stained for DNA content analysis with FxCycle Violet, a DNA-selective dye. Finally, cell cycle analysis was performed using a BD FACSCanto II flow cytometer (BD Bioscience). Data was analyzed using FlowJo V10 (FlowJo, LLC) software.

### Chromatin immunoprecipitation followed by sequencing (ChIP-seq) analysis

Chromatin immunoprecipitation (ChIP) was carried out using the ChIP-IT Express Enzymatic Chromatin Immunoprecipitation Kit (Active Motif) according to manufacturer’s instructions. Briefly, 5×10^6^ 4T1 cells were seeded onto 15 cm tissue culture dishes. After 48 h of incubation, cells were fixed with 1% formaldehyde for 10 min at room temperature. Cells were washed with ice-cold PBS and formaldehyde cross-linking was quenched using glycine stop-fix solution. Cells were harvested through scraping and pelleted by centrifugation at 2,500 rpm for 10 min at 4°C. After cell lysis with ice-cold lysis buffer and a dounce homogenizer, chromatin was sheared using an enzymatic shearing cocktail for 12 min at 37°C. The shearing reaction was stopped with 0.5 M EDTA and the chromatin was separated through centrifugation at 15,000 rpm for 10 min at 4°C. 60-70 μg of chromatin were used with 25 μl of Protein G magnetic beads and 2 μg of specific antibodies for each ChIP reaction. The following antibodies were used for the ChIP reactions: anti-rabbit NUP210 (Bethyl Laboratories), anti-rabbit H3K27me3 (Cell Signaling Technology), anti-rabbit H3K4me3 (Millipore), and anti-mouse Nuclear Pore Complex (NPC) antibody (clone mAb414, Abcam). After incubating the reaction mixture overnight at 4°C on a rotator, the beads were washed with ChIP buffers and DNA was eluted with elution buffer. DNA was then reverse-cross linked, proteinase K-treated, and DNA quality was assessed using a Agilent Bioanalyzer before ChIP-seq library preparation. The library was prepared using a TruSeq ChIP Library Preparation Kit (Illumina) and pooled samples were sequenced on the NextSeq platform.

ChIP-seq data was analyzed using the Sicer algorithm^82^ with default parameters. For narrow peaks such as H3K4me3 enrichment, a window size of 200 was used. For broad peaks such as H3K27me3 and NUP210, a window size of 1000 was used. The ChIP-seq plot profile was generated using deepTools2^83^. For ChIP-seq peak annotation, the ChIPSeeker^84^ Bioconductor package was used. BigWig files were displayed using the Integrative Genomics Viewer (IGV)^85^.

### Publicly available ChIP-seq and Hi-C data analysis

H3K27Ac ChIP-seq data from mouse mammary luminal cells were downloaded from Gene Expression Omnibus (GEO) (Accession No: GSE116384)^26^. CTCF ChIP-seq data from mouse mammary epithelial cells were obtained from GEO (Accession No: GSE92587, sample: GSM2433042)^20^. CTCF and H3K27Ac ChIP-seq data from the MCF7 human breast cancer cell line were obtained from GEO (Accession No: GSE130852). Publicly available Hi-C data from mouse embryonic stem cells were derived from Bonev et al.^37^ and displayed using the 3D Genome Browser^38^.

### RNA isolation and quantitative reverse transcriptase real-time PCR (qRT-PCR)

Cells were directly lysed on cell cultures plate with 1 ml TriPure Isolation Reagent (Sigma). After adding 200 μl chloroform and centrifuging at 14,000 rpm for 15 min at 4°C, the upper aqueous layer containing the RNA was transferred to new tube. To precipitate RNA, 500 μl of isopropanol was added to each tube, which was then vortexted and incubated at −20°C for 1 h. The RNA was further purified using the RNeasy Mini Kit (Qiagen) with on-column DNase (Qiagen) digestion according to manufacturer’s instructions. 2 μg of total RNA was used for cDNA preparation using the iScript cDNA synthesis kit (Bio-Rad). A 1/10 dilution of cDNA was used for qRT-PCR analysis using the FastStart Universal SYBR Green Master Mix (Roche). Sequences of the primers used are listed in the reagents or resource table.

### ChIP qPCR analysis

Chromatin was isolated from 4T1 and 6DT1 cells using the ChIP-IT Express Enzymatic kit (Active Motif) according to manufacturers’s instructions as mentioned above. 10 μl of normal rabbit IgG (Cell Signaling Technology), 10 μl of rabbit CTCF (Cell Signaling Technology), and 10 μl of rabbit H3K27Ac (Cell Signaling Technology) antibodies were used for the immunoprecipitation reaction. ChIP DNA was then subjected to qPCR analysis. The percent (%) input method was used for the calculation of CTCF and H3K27Ac enrichment.

### RNA-seq analysis

4T1 cells were seeded onto 6 cm tissue culture dishes at a seeding density of 4×10^5^ cells/dish. After 48 h of incubation, total RNA was isolated using the protocol described above. On-column DNase treatment was performed to eliminate DNA contamination. Library preparation was performed using the TruSeq Stranded mRNA Library Prep Kit (Illumina) and pooled samples were sequenced on a HiSeq2500 with TruSeq V4 chemistry (Illumina). Differential gene expression analysis from RNA-seq data was performed using Partek Flow software. Gene ontology (GO) enrichment analysis of differentially expressed genes was performed using the PANTHER classification system^86^.

### Promoter luciferase assay

24-well tissue culture plates were coated overnight with 100 μg/ml type I collagen. 293FT cells were seeded in antibiotic free medium at a seeding density of 75,000 cells/well of 24-well plate. 250 ng of pGL4.23 *Nup210* luciferase promoter vectors and 25 ng of pRL-TK renilla luciferase vectors were co-transfected using NanoJuice transfection reagent (Millipore). 24 h after transfection, the luciferase assay was performed using Dual-Luciferase Reporter Assay System kit (Promega) according to the manufacturer’s instructions. Luciferase activity was measured using a GloMax 96 Microplate Luminometer (Promega). Firefly luciferase activity was normalized with Renilla luciferase activity. Eight biological replicates were used per condition.

### Cytokine array

4T1 *Nup210* knockdown cells were cultured in low serum condition (1% FBS) in 6-well plates at a seeding density of 500,000 cells per well. 24 h after incubation, cell culture supernatants were harvested and centrifuged to remove the dead cells. 1 ml of supernatant was used for cytokine profiling using the Proteome Profiler Mouse XL Cytokine Array (R & D Biosystems) according to manufacturer’s instructions. Chemiluminescent signals were quantified using ImageJ (NIH) software.

### 3D DNA FISH analysis

For DNA FISH analysis, cells were grown on Lab-Tek chamber slides. Cells were briefly washed with PBS and fixed with 4% paraformaldehyde. Cells were permeabilized with 0.2% Triton X-100 in PBS and then hybridized with labeled probe. For probe generation, BAC clones were purchased from CHORI (https://bacpacresources.org/). The following BAC clones were used for the experiment: *Cxcl* region: clone RP23-374O6; *Postn* region: clone RP23-480C1; *Ccl2* region: clone RP23-99N1, and *Itgb2* region: clone RP23-166E21. All the clones were expanded and DNA was isolated using the FosmidMax DNA Purification Kit (Epicentre). DNA from each clone was labeled through nick translation with either the Atto550 NT Labeling Kit (Jena Bioscience) or the Digoxigenin NT Labeling Kit (Jena Bioscience). Hybridization was carried out in a humidified chamber at 37°C for 16 h. Post-hybridization rapid wash was carried out with 0.4x SSC at 72°C for 4 min. Digoxigenin was detected with a DyLight 594-Labeled Anti-Digoxigenin/Digoxin (DIG) antibody (Vector Laboratories). The slides were stained with DAPI and Z-stack images were captured using a Zeiss LSM880 Airyscan microscope.

### Response to ECM stiffness assay

To test the ECM stiffness effect on tumor cells, polyacrylamide hydrogel-bound cell culture plates with different elastic moduli was used as a mimic of extracellular matrix (ECM) stiffness. 6-well plates with 0.2 kPa (soft) or 12 kPa (stiff) elastic moduli were purchased from Matrigen. Each well of the plate was coated with 20 μg/ml fibronectin (Sigma) in PBS for 1 h at 37°C. After washing the well with PBS, 4T1 cells were seeded on top of soft or stiff matrix at a seeding density of 100,000 cells/well. 48 h after incubation, cells were harvested for RNA or protein isolation.

### Focal adhesion and cell spreading assay

For focal adhesion immunostaining, Lab-Tek 2-well glass chamber slides (Thermo Fisher Scientific) were coated with 50 μg/ml of collagen type I (Gibco) or 20 μg/ml Fibronectin. Cells were seeded at a seeding density of 20,000-50,000 cells/well. After 6 hours or overnight incubation, cells were fixed in 4% paraformaldehyde for 20 min. Cells were then washed with PBS, permeabilized with PBS + 0.1% Triton X-100 for 5 min, and blocked with PBS + 5% normal goat serum. Cells were incubated with phospho-FAK (Y397) antibody for 1 h. After washing with PBS, cells were incubated with Alexa Fluor 594-conjugated rabbit secondary antibody and Alexa Fluor 488-conjugated phalloidin (F-actin staining) for 1 h. The slides were then washed with PBS and mounted using DAPI (1 μg/ml) or VECTASHIELD with DAPI Mounting Medium (Vector Laboratories). Images were captured using a Zeiss 780/880 confocal microscope.

To check the effect of recombinant CCL2 on cell spreading of *Nup210* KO cells, 50,000 4T1 sg-Ctrl and *Nup210* KO cells were seeded onto Lab-tek glass coverslip coated with type I collagen (50 μg/ml). Cells were allowed to attach on coverslip for 1 h and then cells were incubated with murine recombinant CCL2 (Peprotech) for 5 h. Images were captured using a brightfield microscope and cell area was quantified using ImageJ (NIH) software.

### Actin polymerization inhibition using cytochalasin D

For the cytochalasin D treatment of 4T1 *Nup210* knockdown cells, 20,000 cells were seed onto 4-well μ-Slides with polymer coverslips (Ibidi). 24 h later, cells were treated with 1 μM cytochalasin D, a potent inhibitor actin polymerization, for 2 h. After the treatment, cells were washed with complete medium three times and incubated for another 2 h in a 37°C CO_2_ incubator for the recovery of actin polymerization. Cells were then fixed with 4% paraformaldehyde for 30 min and stained with Alexa Fluor 488-conjugated phalloidin for immunofluorescence imaging. Nuclei were stained with 1 μg/ml Hoechst 33342 (Thermo Fisher).

For the western blot assay, 2.5×10^6^ 4T1 cells were seeded onto 15cm cell culture dishes. 24h later, cells were treated with 0.5 μM Cytochalasin D and incubated 24h before harvesting the cells for protein isolation. For RNA isolation, 3×10^5^ cells were seeded onto 6cm dishes. 24h later, cells were treated with 0.5 μM Cytochalasin D and incubated 24h before lysing the cells for RNA isolation.

### Histone modifying enzyme inhibitor treatment

For immunofluorescence microscopy, 4T1 sg-Ctrl and *Nup210* KO cells were seeded onto 4-well μ-Slides at a seeding density of 25,000 cells/well. 24 h later, cells were treated with DMSO, 5 μM EZH2 inhibitor GSK126 (Selleckchem) or 5 μM H3K27me3 demethylase inhibitor GSKJ4 (Selleckchem) for another 24 h. Cells were then fixed with −20°C methanol for 2 min. Histone H3.1/3.2 and H3K27me3 antibodies were used for immunofluorescence staining according to the protocol described above ^28^.

For the qRT-PCR analysis, 2×10^5^ 4T1 *Nup210* KO cells were seeded onto 6 cm dishes. 24 h later, 5 μM GSK126 or 5 μM GSKJ4 was applied to the cells that were then incubated for another 24-48 h. Cells were then lysed, RNA was isolated, and qRT-PCR was performed as described above.

### Live cell imaging assay

For the automated random cell migration assay, 4T1 cells were seeded onto 96-well polystyrene microplates (Corning) at a seeding density of 2,000 cells/well so that the cell density remained sub-confluent until the end of the imaging period. After 24 h of incubation, cells were incubated with complete medium containing 200 ng/ml Hoechst 33342 (Thermo Fisher Scientific) for 1 h. Cells were then transferred to a Nikon Eclipse Ti2 microscope. Images were captured every 12 minutes with a 20x 0.8 NA objective for 24 h. Total light exposure time was kept to 200 milliseconds for each time point. Cells were imaged in a humidified 37°C incubator with 5% CO_2_. Image processing and cell tracking were carried out with custom MATLAB script described previously^87^. Live cell imaging of nuclear size was also performed using the same approach. To observe the effect of GSK126 and JQ drugs on nuclear size, 3,000 cells were seeded onto ibidi 96 well polymer coverslip plates. 24 h later, cells were stained with Hoechst 33342 for 45 min, washed the medium 3 times with PBS and cells were treated with either 5 μM GSK126 or 1 μM JQ1 and imaged for 24 h.

To study the actomyosin tension dynamics of NUP210-depleted cells, Ftractin-mRuby-P2A-mTurquoise-MLC2 reporter^43^, a gift from the Dr. Tobias Meyer of Stanford University (Addgene # 85146), was used. 6DT1 cells were transduced with this reporter and 50,000 cells were seeded onto type I collagen (50 μg/ml) coated Ibidi 4 well glass bottom coverslip. 24 h later, cells were imaged with a Zeiss 880 Airyscan microscope for 1 h (images were taken at every 15 minutes). Cells were then treated with 1 μM Cytochalasin D for 1 h and imaged. After the drug washout, cells were then further imaged (every 15 min) for 15h in a humidified 37°C incubator with 5% CO_2_.

### Cell invasion assay

The cell invasion assay was performed using BioCoat Matrigel Invasion Chambers with 8.0 µm PET Membrane (Corning) according to manufacturer’s instructions. Briefly, 7.5×10^5^ 4T1 and 6DT1 cells were seeded onto the top well containing DMEM with 0.5% serum. In the bottom well, DMEM with 10% serum was used as a chemoattractant. Cells were incubated in a 37°C incubator for 24 h and 48 h for 6DT1 and 4T1 cells, respectively. After incubation, non-invaded cells were removed from the top well using cotton tips. Cells that had invaded into the Matrigel were then fixed with methanol and stained with 0.05% crystal violet. Matrigel membranes containing invaded cells were then cut and mounted onto glass slides with Vectashield mounting medium (Vector Laboratories). Images of entire membranes were captured as segments and then stitched using an EVOS FL Auto 2 microscope (Invitrogen). Image analysis was performed using Image-Pro Premier 3D (Media Cybernetics) software. Percent cell invasion was calculated using crystal violet intensity per membrane area.

### Circulating tumor cell (CTC) analysis

100,000 6DT1 cells with or without *Nup210* knockdown were injected into the fourth mammary fat pad of FVB/NJ mice. Ten mice were used in each group and three mice were kept uninjected for use as healthy controls. One month after injection, mice were anesthetized with avertin injection. Through cardiac puncture, 600-1000 μl blood per mouse was collected in 50 μl of 0.5 M EDTA solution. An equal volume of blood was taken for red blood cell lysis using ACK lysis buffer. 100 μl of the peripheral blood lymphocyte (PBL) fraction was subjected to fixation with 2% paraformaldehyde for 15 min at room temperature. Cells were permeabilized with PBS containing 0.1% Triton X-100. Cells were vortexed briefly and kept at room temperature for 30 min. 0.5% BSA in PBS was added and cells were pelleted by centrifugation. Cells were then resuspended in ice-cold 50% methanol in PBS and incubated for 10 min on ice. 150,000 fixed cells were stained for CD45, a pan-lymphocyte marker, and pan-keratin, a tumor cell marker. Before staining with antibodies, cells were incubated with FcR Blocking Reagent (1:10 dilution; Miltenyi) for 10 min at 4°C. Cells were then stained with APC-conjugated CD45 (1:25 dilution; Miltenyi) and Alexa Fluor 488-conjugated pan-keratin (1:25 dilution, Cell Signaling Technology) antibodies for 10 min at 4°C. After washing with MACS buffer (PBS, 0.5% BSA, and 2 mM EDTA), cells were incubated with 1 μg/ml Hoechst 33342 (Thermo Fisher Scientific) for 5 min. Cells were then washed again with MACS buffer and resuspended in 200 μl buffer for analysis using a BD FACSCanto II flow cytometer. A CTC (CD45-/Cytokeratin+) gate was created based on the staining pattern of 6DT1 tumor cells in culture and primary tumor cells derived from 6DT1-injected mice. Flow cytometry data was analyzed using FlowJo V10 software.

### Immunohistochemistry analysis

Normal human mammary gland tissue section (HuFPT127) and human breast cancer tissue microarray (TMA) slide (BR10010e) were purchased from US Biomax Inc (Rockville). Formalin-fixed, paraffin embedded TMA slides were processed for antigen retrieval for 15 minutes performed using TAR buffer pH 6 (Dako) in a steamer. Slides were washed followed by endogenous peroxidase (3% hydrogen peroxide in methanol) for 20 minutes. Protein blocking was performed for 20 min (Dako). NUP210 primary antibody (Atlas) was applied at a dilution of 1:100 in antibody diluent (Dako) for 1 hour at room temperature. After incubation, slides were washed in TBS-tween, and Evision+System-HRP labelled polymer anti-Rabbit (Dako) was applied for 30 minutes at room temperature. Next, slides were washed in TBST, then signal was developed using 3,3*′*-diaminobenzidine tetrahydrochloride as the chromogen substrate (Vector Lab). Slides were counterstained with hematoxylin. A digital image data file was created for TMA slide by optically scanning with the Aperio AT2 scanner (Leica Biosystems) at 0.25um/pixel resolution (40x objective). HALO (Indica Labs) Artificial Intelligence (AI) neural network software (mini-net, Indica) was utilized to produce a classification algorithm for tumor and stroma area segmentation and quantification. To quantify the NUP210 signal (pixels) in the nuclear envelope of AI-classified tumor cells based upon color and constant image intensity thresholding, a second algorithm was created using HALO Area Quantification to deconvolve co-localization of DAB chromogen and hematoxylin. Once established, the parameters for image processing were held constant across the experiment. Derived labeling ratios were indices produced by dividing immunolabeled areas of nuclear envelope by tumor area (X100).

### Patient datasets analysis

Distant metastasis-free survival analysis on gene expression signature was performed using GOBO tool (http://co.bmc.lu.se/gobo/gobo.pl)^88^. Data on *NUP210* and Histone H3.1 (*HIST1H3A, HIST1H3B*) amplification in human breast cancer patients was derived from Molecular Taxonomy of Breast Cancer International Consortium (METABRIC) data available on cBioPortal (https://www.cbioportal.org/)^89^. METABRIC NUP210 mRNA expression data was also downloaded from the cBioPortal database and manually processed for further analysis. Distant metastasis-free survival data (*NUP210* mRNA) was obtained from the Km-plotter database (https://kmplot.com/analysis/). The JetSet best probe 213947_s_at was used for *NUP210* expression and patients were separated by upper and lower quartile values. Distant metastatis-free survival data (NUP210 protein) was obtained from Km-plotter database protein module. *NUP210* gene expression data on breast cancer metastatic sites, prostate cancer and melanoma were downloaded from Gene Expression Omnibus (GEO).

## QUANTIFICATION AND STATISTICAL ANALYSIS

### Statistical analysis

In case of the *in vivo* animal studies, p-values were calculated using the Mann-Whitney test in GraphPad Prism 8 software and the results were reported as mean ± standard deviation. For qRT-PCR result analysis, p-values were calculated using a two-tailed, unpaired t-test in GraphPad Prism and results were reported as mean ± standard error of mean (s.e.m).

### Microscopy image analysis and quantification

The periphery/total nuclear area intensity ratio of histone H3.1/3.2, H3K27me3, and H3K9me3 was quantified using Fiji software^81^. Briefly, total nuclear area was defined by DAPI staining, while the nuclear periphery was defined as the region 25% towards the interior from the nuclear edge (average distance for sg-Ctrl: 0.97 μm and KO-N13: 0.89 μm). Mean fluorescence intensity was quantified in both the periphery and total nuclear area and the periphery/total ratio was plotted. The heterochromatic foci/total nuclear area intensity of histone H3.1/3.2 was quantified using Fiji software. Heterochromatic foci regions were segmented using DAPI staining. Mean fluorescence intensity was quantified in both heterochromatic foci and total nuclear area and the heterochromatic foci/total ratio was plotted as box plots (bottom and top of the box denote first and third quartile respectively, whiskers denote -/+ 1.5 interquartile range (IQR), horizontal lines denote median values). p-values were calculated using a two-tailed Mann-Whitney U-test.

The position of DNA FISH spots relative to the nuclear centroid and periphery was quantified using the Cell module within the Imaris image analysis suite (Bitplane). To account for differences in nuclear size, the relative position was calculated by dividing the shortest distance of the FISH spot to the nuclear centroid, by the length of the three-point line encompassing the nuclear centroid, FISH spot, and nuclear periphery. Relative positions were plotted as cumulative distribution frequencies using RStudio software. p-values were calculated using a two-tailed Mann-Whitney U-test. The distance of DNA FISH spots to heterochromatic foci was quantified using Fiji software with the 3D region of interest (ROI) manager plugin, TANGO ^90^. Heterochromatic foci were segmented using DAPI staining and used to create a distance map. DNA FISH spots were segmented and the distance to the nearest heterochromatic foci was calculated and plotted as box plots (bottom and top of the box denote first and third quartile respectively, whiskers denote -/+ 1.5 interquartile range (IQR), horizontal lines denote median values). p-values were calculated using a two-tailed Mann-Whitney U test.

Focal adhesion counts per cell were quantified using Fiji software. Cytoplasmic areas were segmented using phalloidin staining. pFAK Y397 stained focal adhesions were segmented as follows. Adhesions were filtered from image noise using a Difference of Gaussian band-pass filter. The filtered image was subsequently segmented using an intensity threshold. Segmented adhesions were counted and quantified on a per cell basis.

Three dimensional surface reconstruction of Z-stack images has been performed using Imaris image analysis suite version 9.7.1 (Bitplane).

## DATA AND CODE AVAILABILITY

Gene expression (RNA-seq) and ChIP-seq data have been deposited in Gene Expression Omnibus (GEO) database under the accession number GSE146591.

